# Current state, existing challenges, and promising progress for *de novo* sequencing and assembly of monoclonal antibodies

**DOI:** 10.1101/2022.07.21.500409

**Authors:** Denis Beslic, Georg Tscheuschner, Bernhard Y. Renard, Michael G. Weller, Thilo Muth

## Abstract

Monoclonal antibodies (mAbs) are biotechnologically produced proteins with various applications in research, therapeutics, and diagnostics. Their ability to recognize and bind to specific molecule structures makes them essential research tools and therapeutic agents. Sequence information of antibodies is helpful for understanding antibody-antigen interactions and ensuring their affinity and specificity. *De novo* protein sequencing based on mass spectrometry is a useful method to obtain the amino acid sequence of peptides and proteins without *a priori* knowledge. Deep learning-based approaches have been developed and applied more frequently to increase the accuracy of *de novo* sequencing. In this study, we evaluated five recently developed *de novo* sequencing algorithms (Novor, pNovo 3, DeepNovo, SMSNet, and PointNovo) in their ability to identify and assemble antibody sequences. The deep learning-based tools PointNovo and SMSNet showed an increased peptide recall across different enzymes and datasets compared to spectrum-graph-based approaches. We evaluated different error types of *de novo* peptide sequencing tools and their performance for different numbers of missing cleavage sites, noisy spectra, and peptides of various lengths. We achieved a sequence coverage of 93.15% to 99.07% on the light chains of three different antibody datasets using the de Bruijn assembler ALPS and the predictions from PointNovo. However, low sequence coverage and accuracy on the heavy chains demonstrate that complete *de novo* protein sequencing remains a challenging issue in proteomics that requires improved *de novo* error correction, alternative digestion strategies, and hybrid approaches such as homology search to achieve high accuracy on long protein sequences.

## 1 Introduction

Monoclonal antibodies (mAbs) are immunoglobulins of unique specificity generated artificially in laboratories to mimic antibodies produced by the immune system (1). Their reproducible production under certain conditions and high binding affinity to target molecules make them essential to various diagnostic and analytical applications in immunology, clinical chemistry, food chemistry, environmental analysis, biochemistry, therapeutics and medicine (2–4). Recently, multiple authors pointed out how antibodies can be the cause of critical quality issues, thereby causing a so-called reproducibility crisis. In fact, the results of multiple landmark papers could not be replicated because monoclonal antibodies often lacked crucial quality control steps (5,6). Several criteria for validation and characterization have been recommended to avoid irreproducibility in research applications. One essential step for improving the research quality includes the confirmation of the amino acid sequence (7). In addition, retrieving sequence information of antibodies is crucial for understanding the structural basis of antibody-antigen binding, recognition, and interaction. The structural basis for the specificity in protein-protein interactions lies in the sequence diversity of antibodies. The majority of sequence diversity focuses on the hypervariable loops within the variable regions of antibodies, called complementarity determining regions (CDRs), which are mainly responsible for the interaction between the antibody and their target structures (8,9). Most established methods for antibody *de novo* sequencing rely on sequencing mRNA from hybridoma cells. However, these approaches all depend on the availability of pure clones of antibody-producing cells (10). Moreover, important post-translational modifications (PTMs), which are impacting the antigen binding, developability, and effector functions, cannot be detected by DNA sequencing (3). Hence, approaches to sequence the antibody on protein level are necessary.

Tandem mass spectrometry (MS/MS) is a powerful method for retrieving the amino acid sequence of peptides. Typically, in standard shotgun proteomics, protein samples are digested with proteolytic enzymes into shorter peptides, which are more suitable for analysis by MS/MS. Conventionally, the peptides of experimentally derived MS/MS spectra are identified by database search, where they are compared to theoretical spectra derived from a protein database. Nonetheless, such an approach depends on the availability of a reference dataset, which is not assured for so far uncharacterized proteins with unique sequences such as antibodies. To obtain sequential information from novel or unknown proteins, *de novo* peptide sequencing is commonly used, which identifies peptides directly from MS/MS spectra without relying on a sequence database. Here, each amino acid is derived by computing mass differences of ions from a fragmented peptide. Since the manual characterization of peptides using *de novo* sequencing can be very time-consuming and difficult, various algorithms have been developed to differentiate signal ion peaks from noise peaks with the goal of predicting the correct peptide sequence (11–13). Most *de novo* sequencing algorithms depend on creating a so-called spectrum graph, which can manage the combinatorial possibilities of peptide predictions from MS/MS data. The basic concept is to represent the information from a mass spectrum by a spectrum graph, where the vertices of the spectrum graph correspond to the peaks of the spectrum. Vertices are connected via an edge if the mass difference between them corresponds to the mass of one or multiple amino acids. Finding a path in the spectrum graph provides a method to generate potential peptide sequences from a mass spectrum (14). Subsequently, various algorithmic methods were published to find the optimal path in the spectrum graph, including machine learning (15), dynamic programming (16) and hidden Markov models (17). Recent advances in deep learning (DL) have marked an important milestone for database-independent prediction of peptide sequences from MS/MS data (13). The encoder-decoder architecture was designed to solve specific tasks in sequence-to-sequence learning. In mass spectrometry, convolutional neural networks (CNNs) are used to encode the mass spectrum, while recurrent neural networks (RNNs) are employed as the decoder to predict the amino acids of the peptide sequence one by one. Multiple methods have been published based on the deep learning paradigm, namely DeepNovo (18), DeepNovo-DIA (19), SMSNet (20), and PointNovo (21).

Although peptide *de novo* sequencing has improved in recent years, the full-length assembly of protein sequences poses another challenging task. In most cases, database algorithms infer the correct proteins from identified peptide sequences. However, the determination of protein sequences, which are not part of public databases, limits the feasibility of this approach. Currently, only few developed methods were reported for a database-independent full-length protein *de novo* sequencing and assembly, for instance, meta-SPS (22), ALPS (23), pTa (24), and MuCS (25). The efficiency of meta-SPS was demonstrated on a protein mixture using multiple fragmentation methods, namely collision-induced dissociation (CID) and higher-energy C-trap dissociation (HCD). Meta-SPS utilized overlapping fragment ion peaks from different spectra to construct meta-contigs before *de novo* sequencing. Across six diverse proteins and the aBTLA antibody, the authors observed a sequence coverage between 68% and 99%. Nonetheless, meta-SPS faces multiple limitations and is only optimized for small mixtures of unrelated proteins (22). Tran *et al*. analyzed antibodies using PEAKS *de novo* (26), PEAKS DB (27) and the homology software SPIDER (28) in a complementary way. The results from these three algorithms serve as an input for their de Bruijn assembler ALPS. On two different antibodies, they achieved a sequence coverage from 96.6 to 100%. Still, despite using homology and database search algorithms, the authors inspected a fragmented and incomplete assembly of long proteins, particularly at the variable region of the heavy chain (23). Thus, *de novo* sequencing of proteins remains a challenging and important problem to date.

Most publications regarding new *de novo* sequencing approaches include a performance comparison of recently developed tools (12,14,29), yet, to our knowledge, there is no published independent evaluation of different *de novo* sequencing algorithms on antibody datasets. In this study, we present a performance evaluation of five recently developed *de novo* sequencing algorithms (Novor, pNovo 3, DeepNovo, SMSNet, and PointNovo), which we chose based on their popularity and availability (Table 2). To compare the ability of previously mentioned tools to reconstruct full length protein sequences without additional database algorithms, we employed the de Bruijn assembler ALPS. Hence, we investigated common error types, the impact of noisy spectra and missing fragmentation ions, and discussed possible solutions and demanding challenges of *de novo* antibody sequencing.

## 2 Materials and methods

### 2.1 Antibody datasets

We evaluated the *de novo* sequencing methods in this study on three publicly available antibody MS/MS datasets, which we describe in the following. We evaluated the Herceptin (Trastuzumab) data set published by Peng *et al*. (10), WIgG1-Mouse, a mAb Mass Check Standard from Waters, and IgG1-Human, a purified human antibody sample from SIGMA-Aldrich, which were both made publicly available by Tran *et al*. (23). *The* datasets were downloaded from the proteomics MS data repositories PRIDE (30) and MassIVE (31). Table 1 gives an overviewofour three evaluated antibody datasets, showing the reference, available digestion, mass instrument, ionization type, and fragment ion resolution. Contrary to WIgG1-Mouse and IgG1-Human, the light and heavy chains of the Herceptin antibody data set were not provided separately. Due to corrupted data files, we could not include the Glu-C digestion of the light chain of IgG1-Human and the trypsin digestion of the light chain of WIgG1-Mouse in our analysis. We therefore analyzed 25 different MS/MS experiments (Supplementary Figure S1).

**Table 1.** Overview of evaluated antibody datasets. For each dataset, we provided the name of the dataset, the ID, the mass instrument, the ionization type, the fragment ion resolution in FWHM, a reference, and the number of proteolytic enzymes in the dataset.

| Dataset Name | Database ID | Mass instrument | Ionization type | Resolution | Ref. | Enzymes |
| --- | --- | --- | --- | --- | --- | --- |
| IgG1-Human | MSV000079<br>801 | LTQ<br>Orbitrap | HCD | 17,500 | (23) | trypsin, chymotrypsin, asp-N, lys-C, glu-C, proteinase K |
| WlgG1-Mouse | MSV000079<br>801 | LTQ<br>Orbitrap | HCD | 17,500 | (23) | trypsin, asp-N, chymotrypsin |
| Herceptin | PXD023419 | Orbitrap<br>Fusion | stepped<br>HCD &<br>EThcD | 30,000 | (10) | trypsin, thermolysin, lys-N, lys-C, glu-C, asp-N, aLP, chymotrypsin, elastase |

### 2.2 Data processing

#### 2.2.1 *De novo* sequencing methods

The MS/MS data files from the previously mentioned datasets were reformatted to Mascot Generic Format (MGF) using ProteoWizard (32) to be compatible with all *de novo* sequencing tools. Table 2 provides an overview of the evaluated *de novo* peptide sequencing algorithms. Further, it provides information about the algorithmic paradigm, the project website, the corresponding reference, and the number of citations as a measure for the popularity.

**Table 2.**
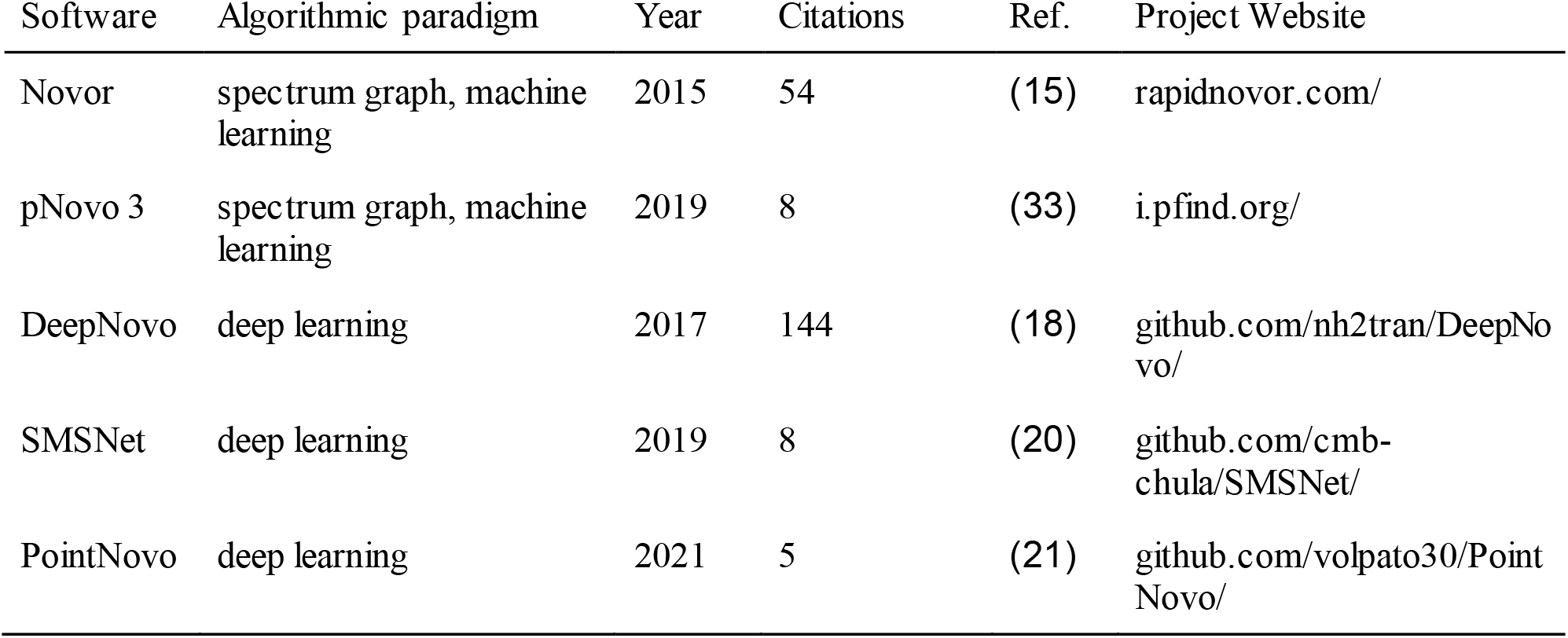
Overview of all *de novo* sequencing tools used in this study. For each algorithm, the name of the tool, the algorithmic paradigm, the year of the publication, the number of citations (as of July 11th), the reference, and the project website of the corresponding method are displayed.

| Software | Algorithmic paradigm | Year | Citations | Ref. | Project Website |
| --- | --- | --- | --- | --- | --- |
| Novor | spectrum graph, machine learning | 2015 | 54 | (15) | rapidnovor.com/ |
| pNovo 3 | spectrum graph, machine learning | 2019 | 8 | (33) | i.pfind.org/ |
| DeepNovo | deep learning | 2017 | 144 | (18) | github.com/nh2tran/DeepNovo/ |
| SMSNet | deep learning | 2019 | 8 | (20) | github.com/cmb-chula/SMSNet/ |
| PointNovo | deep learning | 2021 | 5 | (21) | github.com/volpato30/PointNovo/ |

##### Description of *de novo* sequencing algorithms used

Novor (15) is based on a decision tree scoring function to select peptide predictions. pNovo 3 (33) employs a learning-to-rank framework using gap features and predictions from pDeep (34) to improve the scoring of peptide sequences. Novor and pNovo 3 were developed based on the spectrum-graph approach while using extensive machine learning algorithms for an enhanced scoring function. DeepNovo (18) was the first approach to incorporate the encoder-decoder paradigm for *de novo* peptide sequencing. SMSNet (20) uses a similar CNN-and RNN-based framework but additionally includes optional post-processing and a shift layer in the encoder module. PointNovo (21) adopts an order invariant network structure for the prediction process of higher-resolution data.

##### Parameters for *de novo* sequencing algorithms

We executed Novor (v.1.05) via the DeNovoGUI command-line interface (v.1.16.6) (35). We ran pNovo 3 (v.3.1.3) via its executable GUI, which included pre-trained models for specific enzymes. To perform a fair comparison between spectrum-graph-based tools like Novor and pNovo 3, which are released only with pre-trained models, and the DL algorithms, DeepNovo (v.PNAS), SMSNet, and PointNovo (v.0.0.1), we trained all DL-based tools on high-resolution MS/MS data from the human proteome using the HCD library from MassIVE, which consists of 1,114,503 different peptides (36). Training them on specific antibody data would give DL programs an unfair advantage compared to pre-trained software. We split the spectra into training, validation, and test sets at a ratio of 98:1:1 while making sure that the split datasets did not share any common peptides. Each model was trained for 10 epochs using pre-defined parameters from each tool. We executed all tools at a precursor tolerance of 10 ppm and fragment mass tolerance of 0.02 Da. For each algorithm, carbamidomethylation of cysteine (C+57.02 Da) was set as a fixed modification. Oxidation of methionine (M+15.99 Da) and deamidation of asparagine and glutamine (N+0.98 Da & G+0.98 Da) were set as variable modifications. DeNovoGUI and the DL tools were executed on a Linux server machine (100 cores, 64GB RAM). We executed pNovo 3 on a Windows 64-bit computer since the software was not supported for a Linux operating system.

#### 2.2.2 Assembly of identified peptides

The predicted peptides were further processed by the de Bruijn sequence assembler ALPS (23) to evaluate the ability of different *de novo* sequencing tools to reconstruct complete protein sequences. Other assembly tools are either not combinable with recently developed *de novo* sequencing algorithms (meta-SPS (22)) or they are specially designed to perform on a semi-random approach instead of enzyme-specific cleavages (pTa (24) & MuCS(25) for *microwave*-assisted acid hydrolysis). Based on the peptide sequences and confidence scores for each amino acid position, ALPS can assemble contigs from short *de novo* subsequences. As described by the authors, a k-mer size of k = 7 ensures a sufficiently high coverage of the amino acid sequence while preventing repetitiveness of the resulting contigs at the same time. ALPS takes the *de novo* confidence score into consideration for the assembly, but it could generate incorrect results using a high amount of low-confidence k-mers. The authors of DeepNovo recommend removing sequence contaminants from *de novo* sequencing results by excluding peptides with a confidence score below50to improve the quality of the assembly (18). Qiao *et al*. argued that a confidence of 35 can be considered an accurate score (37). Since every single *de novo* sequencing algorithm calculates its confidence score in a different manner, we chose the threshold for the confidence score based on the amino acid-level precision. We removed peptide sequences below the confidence score of each tool for which the AA precision was below 50%. This aims to filter out low-quality predictions and at the same time ensures that we do not miss correctly predicted peptides, which have been assigned a low confidence score by the corresponding tool. We aligned the target contigs with the ground truth antibody sequence to classify the assembly results using BLAST (38). Based on the alignments of the top contigs, we calculated the protein coverage and accuracy. The target sequence was regarded as being covered in case a contig was aligned to the target (sub-)sequence. We calculated the accuracy by the number of correct sequence calls, which were aligned to the target sequence.

### 2.3 Evaluation metrics

#### 2.3.1 Database Search

For validating *de novo* sequencing algorithms, we compared each prediction to a pseudo-ground truth, which is commonly obtained by database search (12,39). Since the evaluated datasets do not include a labeled ground truth for each spectrum, we performed a database search using the antibody sequences as our protein database. We used the combined results of the database algorithms MS-GF+ (40) and X!Tandem (41), which were both executed via SearchGUI (v.4.1.7) (42) and post-processed via PeptideShaker (v.2.2.2) (43) on a 64-bit Windows computer. We filtered all resulting peptide-spectrum matches (PSMs) using a false discovery rate (FDR) of 1%. The combined results of two database algorithms and an FDR rate of 1% would generate a reliable pseudo-ground truth for the evaluation of the *de novo* sequencing tools. We chose the cleavage parameters according to the used enzyme of the provided input file. Furthermore, the search parameters included the same modifications and mass tolerance that we selected for the *de novo* sequencing algorithms.

#### 2.3.2 Recall and precision

We compared the predictions of each *de novo* sequencing algorithm with the pseudo-ground truth peptides, which were identified by database search. Recall and accuracy were measured at the peptide and amino acid level. The recall at the peptide level is defined as follows:

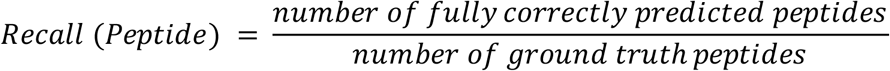

The performance at the amino acid level was measured by matching amino acids between the prediction and the ground truth. We applied the same evaluation metric adapted by DeepNovo, Novor, and PointNovo (15,18,21): amino acids were considered as matched ones if their masses were different by less than 0.1 Da and if the prefix masses before them were different by less than 0.5 Da. The amino acid recall and, respectively, precision are defined as follows:

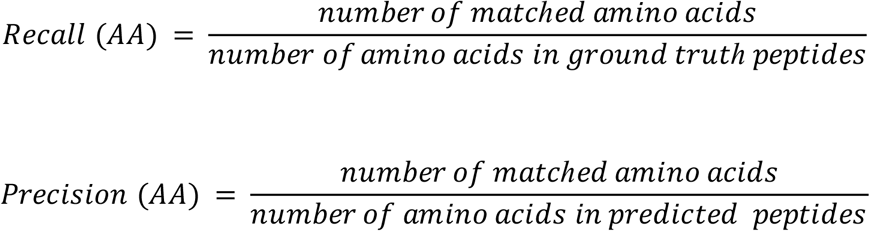

Furthermore, we used the confidence scores of each *de novo* sequencing algorithm to generate precision-recall curves and calculate the area under the curve (AUC), which can be interpreted as a summary of the sequencing accuracy on amino acid level (44).

#### 2.3.3 Identification of fragment ions and noise

To evaluate the amount of noise and missing fragment ions in each spectrum, we labeled each peak as a peptide peak or a noise peak using the Pyteomics framework (45). For each cleavage site we tried to identify 8 different ion types (b, b(2+), b-NH3, b-H2O, y, y(2+), y-NH3, y-H2O), since all evaluated *de novo* sequencing algorithms take these ion types into consideration for retrieving the peptide sequence. If possible, we matched these ion types to corresponding peaks in the spectra within a tolerance of 0.5 Da. Otherwise, we declared the cleavage site as missing. We only considered noise peaks if their intensity exceeded the median noise intensity for each dataset. The number of noise peaks above this threshold were used to calculate the noise factor, which is defined as the ratio of the number of high intensity peaks and the number of fragment ion peaks. McDonnell *et al*. applied this approach recently in their evaluation of *de novo* sequencing algorithms (29).

## 3 Results

### 3.1 Performance of *de novo* sequencing algorithms on antibody data at the peptide and amino acid level

We evaluated five state-of-the-art *de novo* peptide sequencing algorithms, namely Novor, pNovo 3, DeepNovo, SMSNet, and PointNovo. For this purpose, we used the antibody data sets described in section 2.1 to measure the accuracy across different enzymes using metrics specified in section 2.3. Using three antibody datasets, we relied on 183,873 MS/MS scans, from which 23,844 peptides were identified with database search. Peptide identifications by database tools served as a ground truth for evaluating the predictions from *de novo* sequencing tools. By comparing *de novo* sequencing results to this reference, we were able to identify the number of correctly predicted amino acids and peptides for each tool.

Each algorithm generates a confidence score along the predicted sequence to reflect its quality. Setting a threshold to the confidence score outputs different sets of predicted peptides. A high threshold would show a small number of peptides with high precision, but it would exclude a large part of the dataset, consequently reducing the recall. Here, we used different thresholds of the confidence score to draw precision-recall curves and use the area under curve (AUC) as a summary metric for the accuracy of *de novo* sequencing results. Figure 1 displays the precision-recall curves (A-C) and the AUC (D) of *de novo* sequencing tools across six different enzymes of the Herceptin dataset. The five evaluated algorithms show an overall higher AUC on trypsin and thermolysin than on other proteases. The performance is generally lower on the enzymatic datasets of glu-C and lys-N. Lower efficiency of non-tryptic enzymes for the prediction of peptide sequences was reported in different publications and had several reasons (46–48). First, trypsin shows a higher number of peptide-spectrum matches, which is caused by a bias of database search algorithms towards peptides digested with trypsin (49). Furthermore, peptidesdigestedwith trypsin are better suited for HCD fragmentation since they include at least one positive charge at each terminus, generating reliable b- and y-ion fragmentation patterns. In contrast, non-tryptic proteases may lack positive-charged termini, which makes it more challenging to identify the correct peptide (50). The AUC on aLP, asp-N, elastase, chymotrypsin, glu-C, and lys-N is considerably lower across all tools due to their distinct cleavage patterns.

**Figure 1.**
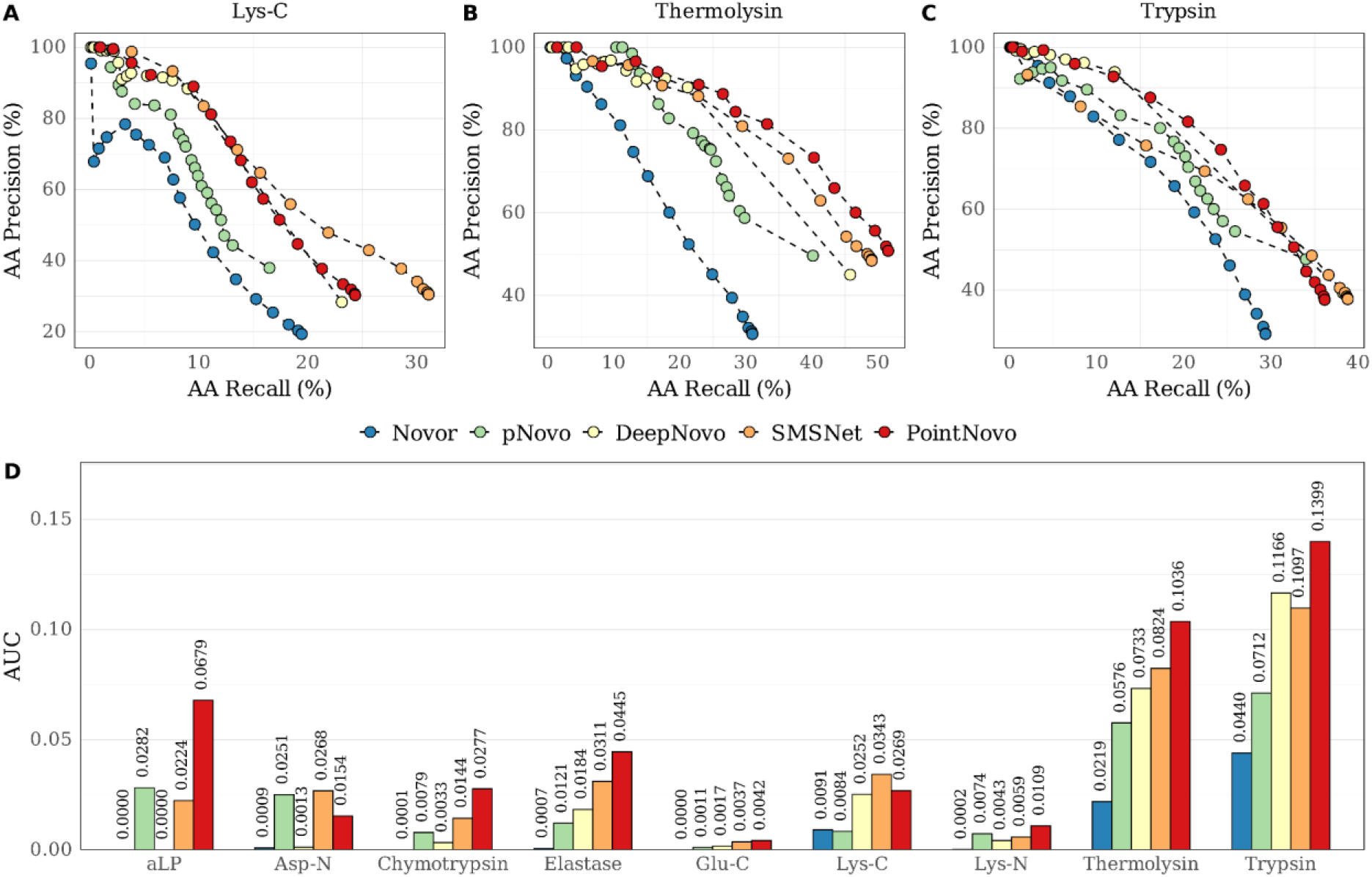
The precision-recall (PR) curves of Novor, pNovo 3, DeepNovo, SMSNet, PointNovo for lys-C (A), thermolysin (B), and trypsin (C) of the Herceptin dataset. The area under curve (AUC) of the five algorithms for each PR curve and each enzyme of Herceptin (D).

Among the established *de novo* sequencing tools, SMSNet shows a higher AUC in relation to PointNovo on lys-C and asp-Ndata. On aLP, chymotrypsin, elastase, glu-C, lys-N, thermolysin and trypsin, PointNovo shows a higher AUC. Novor recalled fewer PSMs on non-tryptic enzymes since its freely available software version is only trained and optimized on tryptic MS/MS data. The high AUC of pNovo 3 on non-tryptic peptides can be attributed to the pre-trained models for specific enzymes. A pre-trained enzyme-specific model was not available for elastase and thermolysin, which explains the low accuracy of pNovo 3 on these datasets. We trained the DL tools on a large dataset, which mainly consisted of tryptic peptides, explaining the overall lower performance on non-tryptic enzymes like asp-N, glu-C, lys-N, and chymotrypsin. We observe comparable results on the IgG1-HC dataset (Supplementary Figure S2), where the DL tools show a higher AUC on the tryptic dataset compared to Novor and pNovo 3. In contrast to the Herceptin dataset, the DL tools show a higher performance on lys-C than on trypsin, which can be explained by the similar cleavage specificities of lys-C and trypsin (51).

Figure 2 displays the total peptide recall (A), amino acid recall (B), and amino acid precision (C) across all nine enzymatic cleavages of Herceptin. In contrast to the results shown in Figure 1, we used all predictions from each tool regardless of their confidence score. Here, either SMSNet or PointNovo show the highest amino acid recall compared to all single *de novo* algorithms across all enzymes (Figure 2B). However, the peptide recall of SMSNet is remarkably lower compared to its relatively high AA recall. While SMSNet shows the highest AA recall on elastase (41.59%), its peptide recall (9.68%) is lower than that of PointNovo. We detect similar differences between the peptide and amino acid recall on different enzymes of the IgG1-HC dataset (Supplementary Figure S2). While examining specific MS/MS spectra, we observed that SMSNet tends to generate unusually long predictions for low-quality mass spectra with incomplete fragmentation patterns. The same observation was also reported by Qiao *et al*. (21), where the authors concluded that the peptide recall is a more reliable metric for comparing the performance of *de novo* sequencing tools. Regarding the recall on peptide level, PointNovo shows a higher number of correct peptide predictions compared to DeepNovo and SMSNet on trypsin, thermolysin, lys-N, glu-C, elastase, chymotrypsin, and aLP. pNovo 3 predicted a similar or even higher number of correct peptides of the non-tryptic datasets asp-N and aLP than the deep learning tools due to its enzyme-specific models. Furthermore, pNovo 3 shows a superior accuracy on amino acid level across all datasets (Figure 2C) due to its extensive reranking process.

**Figure 2.**
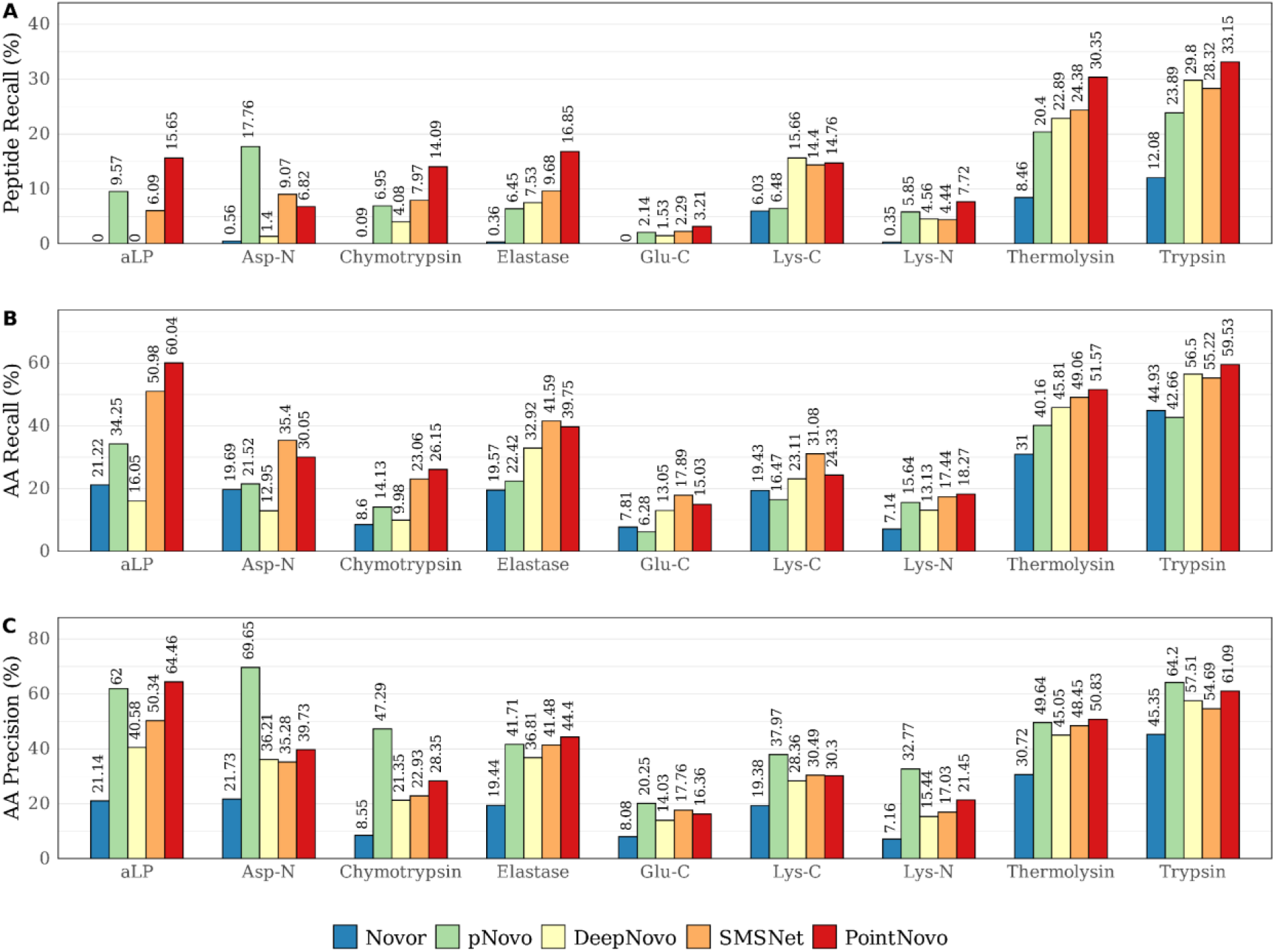
Total recall and precision of Novor, pNovo 3, DeepNovo, SMSNet and PointNovo across different enzymes on Herceptin. (A) Recall at peptide level. (B) Recall at amino acid level. (C) Precision at amino acid level.

Correspondingly, Figure 3 shows the precision and recall values on peptide and amino acid level using only tryptic datasets across all available datasets. Here, the DL tools (DeepNovo, SMSNet, PointNovo) show superior performance compared to spectrum-graph-based approaches. Furthermore, we found overall differences in performance between antibody datasets even when using identical enzymatic approaches and instruments. For example, the peptide recall of PointNovo for tryptic data was 30% higher on the WIgG1-Mouse-HC dataset than on IgG1-Human-HC. SMSNet shows the highest peptide recall (26.42%) on the light chain of IgG1-Human but obtains the lowest peptide recall (10.88%) on the heavy chain of the same antibody.

**Figure 3.**
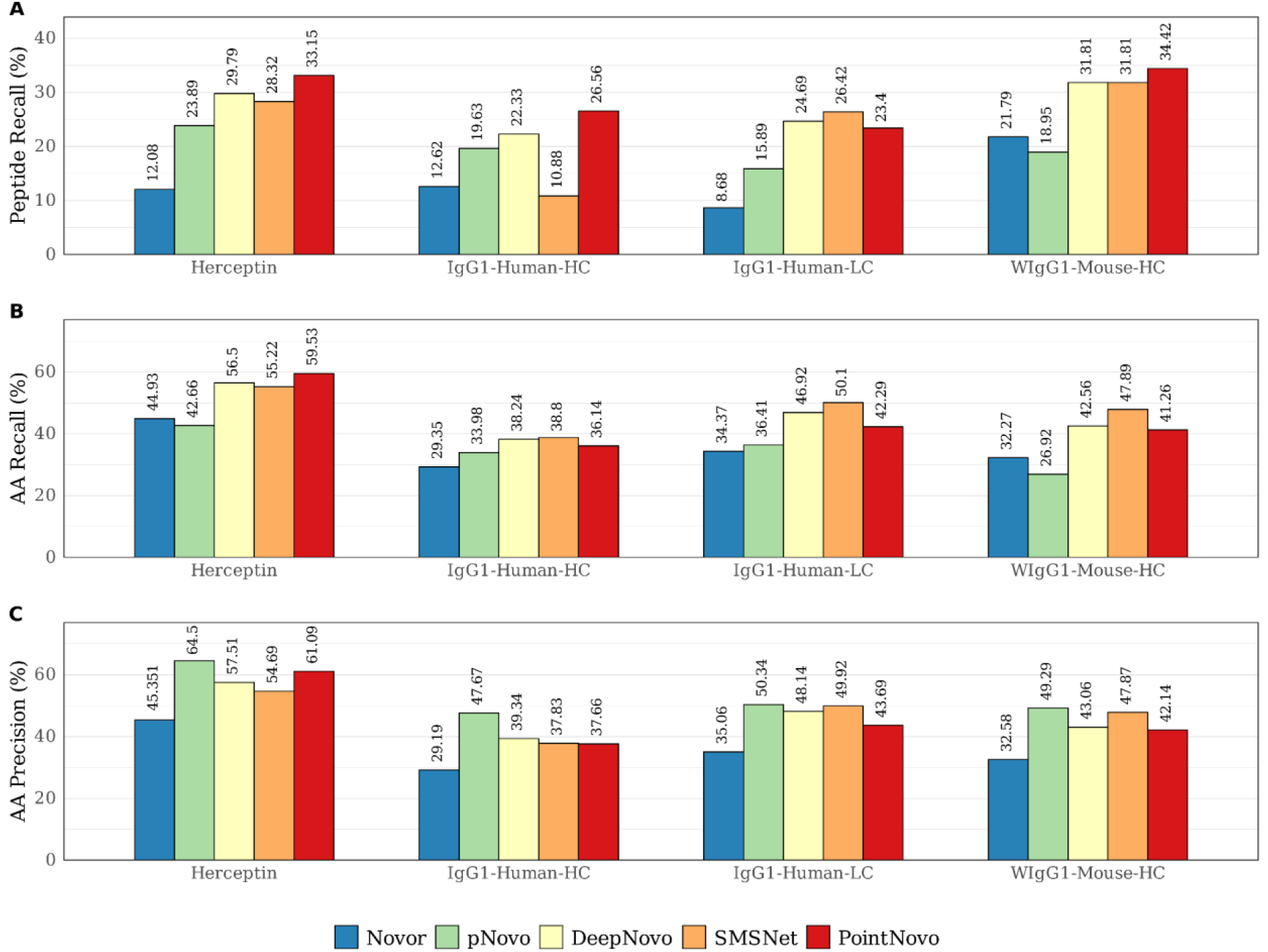
Total recall and precision of Novor, pNovo 3, DeepNovo, SMSNet and PointNovo for trypsin across different datasets. (A) Recall at peptide level. (B) Recall at amino acid level. (C) Precision at amino acid level.

Supplementary Figure S6 shows the number of shared predictions between the different *de novo* sequencing tools for IgG1-Human-HC trypsin. Overall, the tools tend to generate distinct peptide predictions and only share overall few common identifications, which could make a complementary approach useful for re-evaluating peptides sequence candidates.

To evaluate a possible complementary use of multiple algorithms, we compared the number of correctly identified peptides and amino acids for each pair of the five tools for the Herceptin trypsin dataset (Figure 4). Here, the combined results of PointNovo and DeepNovo achieve a peptide recall of 39.60%, whereas PointNovo by itself reaches a peptide recall of 33.15%. At the amino acid level, the joint predictions of PointNovo and SMSNet show an increased recall of 65.11% compared to the single predictions of PointNovo, reaching an AA recall of 60.62%. A similar increase in performance can be observed for the WIgG1 dataset (Supplementary Figure S7), where PointNovo and SMSNet achieve a joint peptide recall of 31.86%, hinting at the potential of a complementary approach.

**Figure 4.**
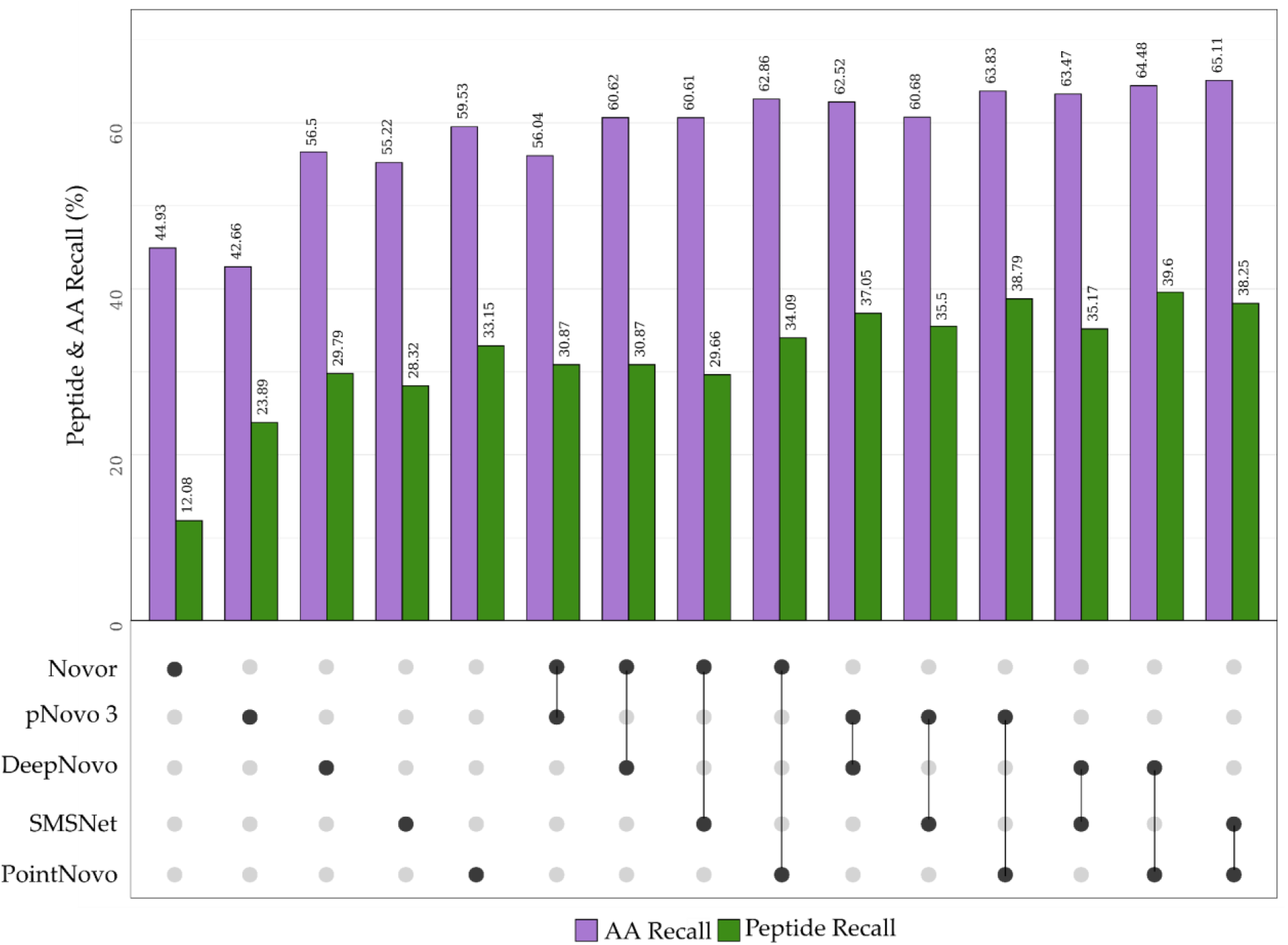
The AA recall (violet) and peptide recall (green) by each pair of combined *de novo* sequencing predictions for the Herceptin trypsin dataset.

### 3.2 Evaluation of error types

Following the performance on peptide and amino acid level, we evaluated the source of incorrect predictions of the different *de novo* sequencing algorithms. McDonell *et al*. reported previously that missing fragment ions and noise peaks pose a challenge for *de novo* sequencing algorithms (29). We observed that 90.51% of all 23,227 validated spectra were missing at least one fragment ion. Furthermore, we detected that 84.32% of all peaks from these spectra were classified as noise peaks. In Figure 5, we show the peptide recall for different numbers of missing cleavage sites and different noise factors of all validated spectra. As expected, *de novo* sequencing algorithms tend to identify a higher number of correct peptides from spectra with a lower amount of missing fragment ions (Figure 5A). Missing fragment ions decrease the overall performance of all *de novo* sequencing tools. PointNovo shows a superior performance on spectra with up to three missing cleavage sites. On spectra with at least four missing cleavage sites, all deep learning-based algorithms show a low peptide recall of 9.56% to 14.62%, while PointNovo and SMSNet perform slightly better than DeepNovo and pNovo 3. Novor shows a noticeable lower performance compared to all other algorithms.

**Figure 5.**
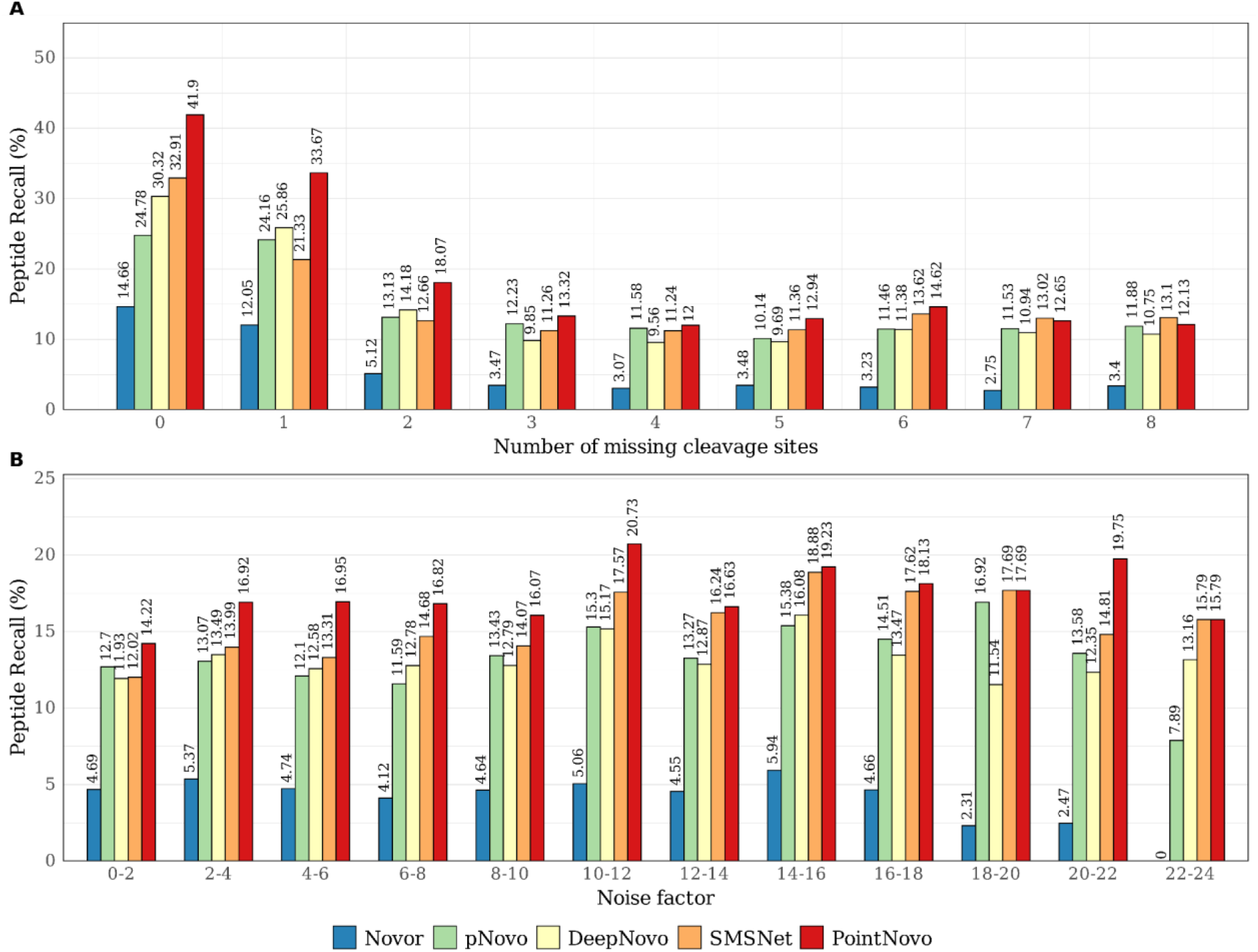
Total peptide recall of Novor, pNovo 3, DeepNovo, SMSNet and PointNovo across all datasets for different number of cleavage sites missing (A) and different noise factors (B) of the specific spectra.

When viewed alone, the noise factor of different spectra does not have a strong effect on the accuracy of the *de novo* sequencing algorithms (Figure 5B). As McDonell stated, this is due to the stronger influence of the number of missing fragmentation sites on the peptide recall of each tool. Supplementary Figure S10 shows the impact of both, the noise factor, and the number of missing cleavages, on the accuracy of PointNovo, SMSNet, and pNovo 3. This demonstrates how a noise factor of at least 2 is already decreasing the prediction accuracy on spectra with no missing cleavage sites across all evaluated tools.

Next, we compared the relation between the peptide length and prediction accuracy. The number of errors increases with a higher peptide length since the probability of a missing ion fragment is more likely (12). We considered peptides of a sequence length between 8 and 20 and summarized peptides with a length of over 20 amino acids. Figure 6 displays the peptide and amino acid recall according to their peptide length from Novor, pNovo 3, DeepNovo, SMSNet and PointNovo. As expected, the number of correct predictions was higher for peptides below a size of 14 amino acids. Tryptic and other proteolytic enzymes generate peptides of this length. Miscleavages lead to peptides of greater length, which would overall lower the prediction accuracy. For length 8 peptides, PointNovo correctly predicts 50.72% of all spectra, while for length 10 peptides it only predicts 26.18% of all spectra correctly. Supplementary Figure S12 shows a similar plot using absolute numbers of correct and incorrect predictions. While PointNovo and SMSNet predict a higher number of correct peptides for a length over 20 compared to pNovo 3, the number of incorrect predictions increases for SMSNet and PointNovo at a greater length. None of the evaluated tools predicted over 100 correct peptides for this length, indicating that even recently developed deep learning tools are not suited for the prediction of long peptides.

**Figure 6.**
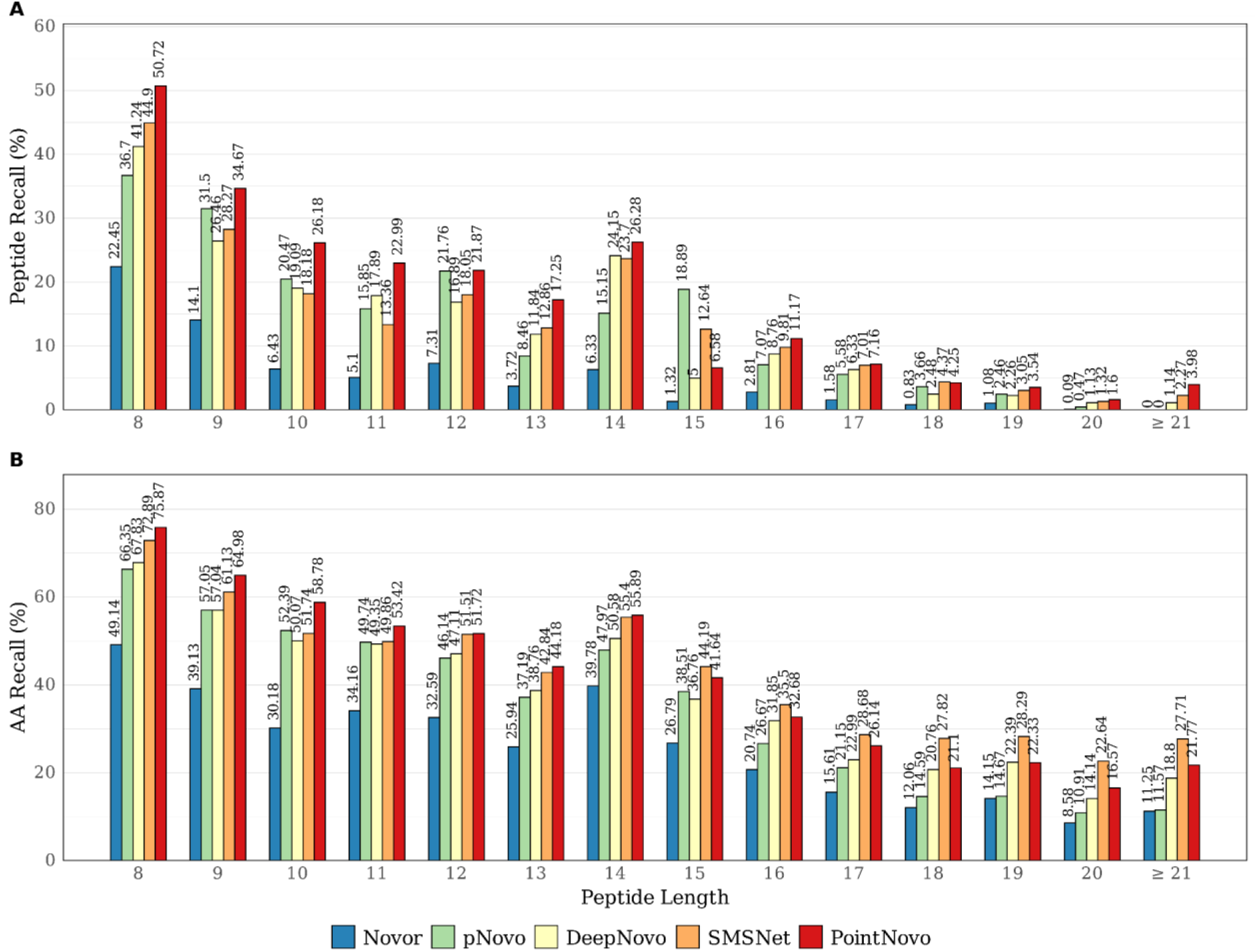
Total peptide recall (A) and AA recall (B) of Novor, pNovo 3, DeepNovo, SMSNet and PointNovo across all datasets for different peptide lengths.

Further, we investigated the relationship between peptide length, number of missing cleavage sites and prediction accuracy (Figure 7). The prediction accuracy decreases from short peptides with few missing fragmentation sites to long peptides with a high number of missing cleaves for each algorithm. The DL-based tools SMSNet and PointNovo show a higher prediction accuracy for peptides of a greater length compared to Novor and pNovo 3. However, even with all cleavage sites present, the evaluated algorithms only rarely identified correct peptides with a length of at least 18 amino acids.

**Figure 7.**
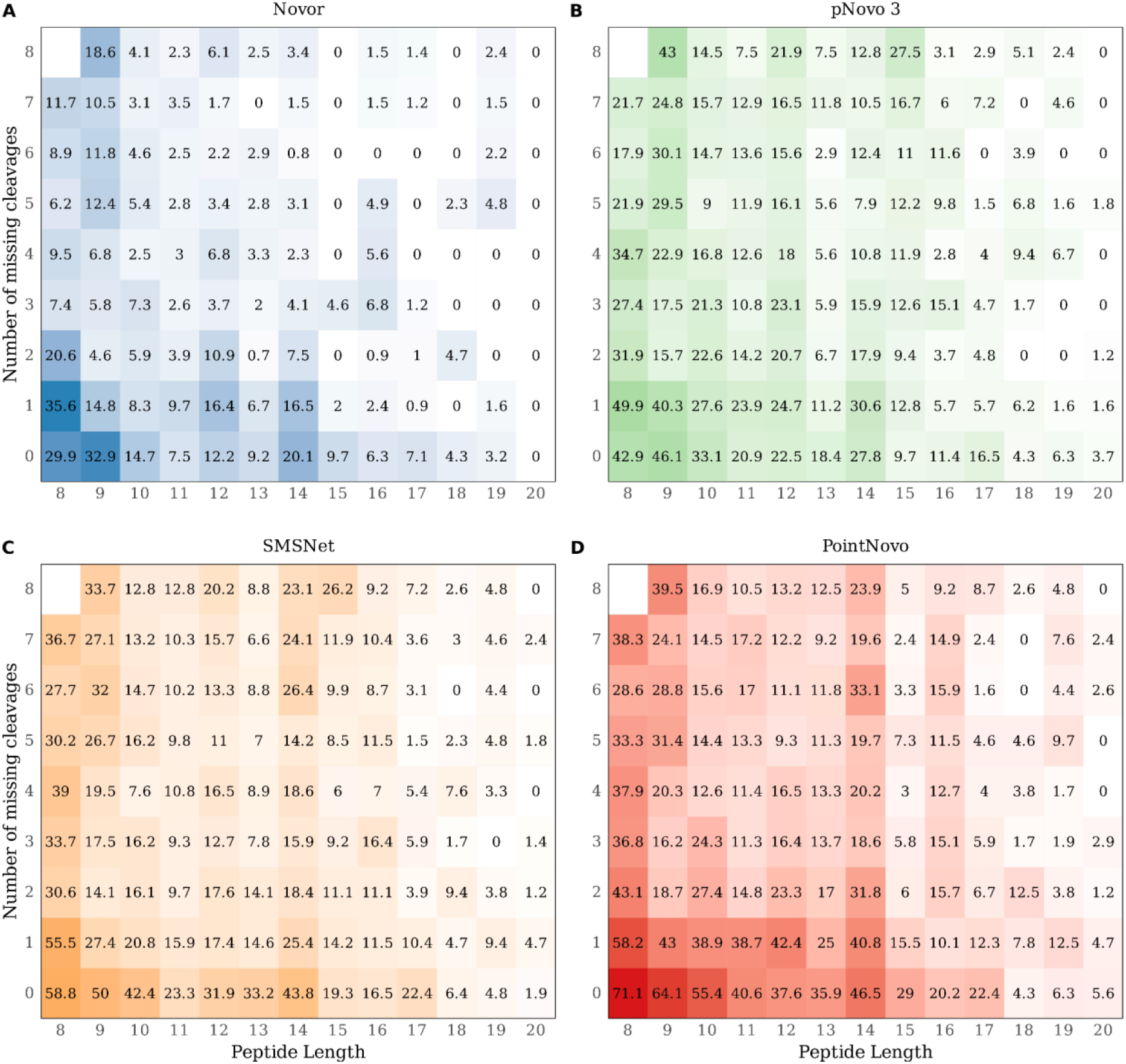
Heatmap showing peptide recall for different number of missing cleavages (y-axis) and peptide lengths (x-axis). Higher peptide recall is shown in blue for Novor (A), green for pNovo 3 (B), orange for SMSNet (C), and red for PointNovo (D). Lower peptide recall is displayed in white. Spectra are not distributed uniformly and the squares on the right and top of the plots include less spectra, since combinations of long peptides and a high number of missing cleavages (top right) occur less likely.

Following the influence of peptide length on the predictive performance, we compared the frequency of certain error types across multiple tools. We categorized incorrect peptide sequence predictions into 11 different error types and compared their relative amount between SMSNet, PointNovo, and pNovo 3 across all datasets (Table 2). We observed that most errors were caused due to more than six wrongly assigned amino acids. Among the error types under 6 AAs, the inversion of the last 3 amino acids (PointNovo) and the replacement of 1 AA by 1 or 2 AAs (SMSNet, pNovo 3) appear as the most frequent origins of incorrect peptide predictions. SMSNet and PointNovo generated more predictions with a higher number of mismatches in relation to pNovo 3 (44.90%). Furthermore, the number of inversions was slightly lower on pNovo3 demonstrating the advantage of a re-ranking framework for improved accuracy.

In Supplementary Table S1, we show the relative amount of amino acid substitutions for predictions, where only 1 amino acid was incorrectly assigned. Here, the misidentification between deamidated asparagine and aspartic acid makes up 41.64% (SMSNet) to 74.51% (PointNovo) of all misidentifications for predictions with at most one sequencing error. Replacements of amino acids can be caused by mass ambiguities since certain amino acids share a similar mass, like lysine and glutamine (mass difference of 0.036 Da). Combinations of different amino acids can result in an identical mass, such as the mass of one glutamine compared to the mass of one alanine and one glycine (no mass difference). Furthermore, posttranslational modifications can influence the residue mass, for instance, the deamidation of asparagine converts it to aspartic acid or isoaspartic acid. Missing fragment ions lead to inversions, where the correct order of amino acids cannot be derived correctly. Additionally, the conflation of y-ions with b-ions can be another reason for inversions (11). Overall, the evaluated *de novo* sequencing tools generated a high number of incorrect peptide predictions due to complete misidentifications. In Supplementary Table S2, we show the error types on spectra without missing cleavage sites. There, pNovo 3, SMSNet and PointNovo show a lower number of complete misidentifications, suggesting that the number of missing cleavage ions pose a limitation to all *de novo* sequencing tools, including algorithms with an extensive re-ranking approach such as pNovo 3.

### 3.3 Database-independent assembly of predicted peptide sequences

To validate the predictions of different peptide *de novo* sequencing tools on assembly level, we used the de Bruijn graph assembler ALPS, which generates several contiguous sequences (contigs) based on the *de novo* peptide results and their confidence scores (23). We compared the longest constructed contig, the overall sequence coverage, and the sequence accuracy for three antibody samples. As described in section 2.2.3, we considered only aligned contigs for the calculation of sequence accuracy. We excluded contigs from the analysis in case they only covered sequence regions, which were already reported by other contigs. We aligned the resulting sequences against the target protein to evaluate the overall coverage and number of mismatches. The light chains are 210 to 219 AAs long, while the heavy chains of our evaluated antibodies include over 440 AAs, which present a challenge for a complete sequence assembly. The longest constructed contig for the heavy chain of WIgG1 was generated by SMSNet, covering only 71 AAs (16.10%) of the protein sequence. On the light chain of WIgG1, the results of PointNovo were concatenated to a contig, which covered 108 AAs (49.32%) of the entire sequence.

Since single contigs only cover a small region of the full-length protein, we evaluated the protein sequence coverage using a higher number of contigs for the light chain of IgG1-Human (Table 4). Combined with PointNovo, we were able to assemble 93.15% (WIgG1) to 99.07% (Herceptin) of the whole antibody sequence with an accuracy of 89.62% to 93.63%. We observed a high sequence coverage for SMSNet from 90.74% (IgG1) to 97.20% (Herceptin). In contrast, pNovo 3 achieved a coverage from 35.62% (WIgG1) to 85.65% (IgG1). Interestingly, we were able to achieve a high coverage and accuracy on the light chain of WIgG1, although we only used the enzymatic datasets of chymotrypsin and asp-N.

**Table 3.**
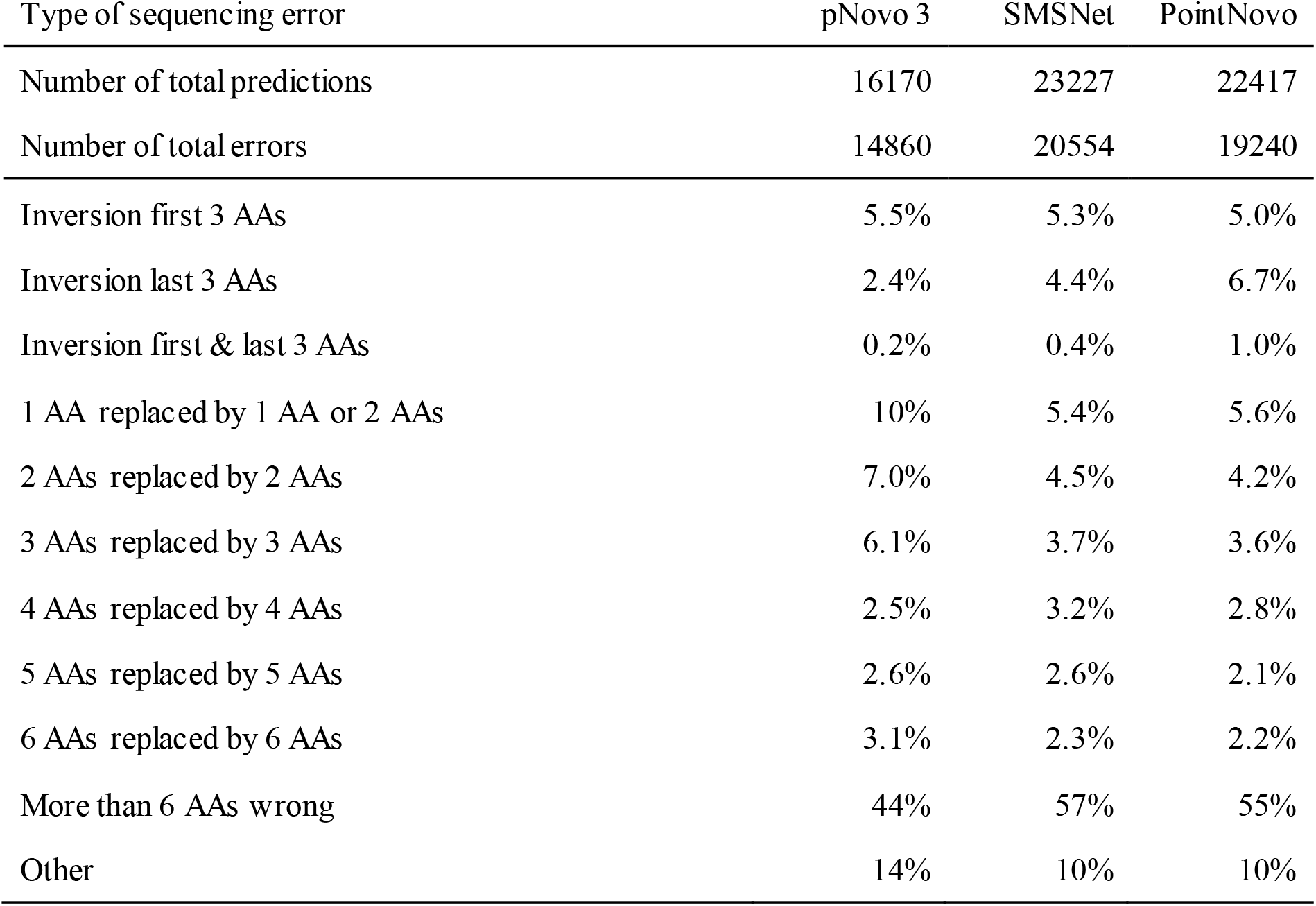
Error types made by *de novo* sequencing algorithms tools pNovo 3, SMSNet, PointNovo on the datasets of IgG1-Human, WIgG1-Mouse, and Herceptin. Shown are the total number of predictions, total number of errors and the relative amount of 11 different error types for each algorithm. ‘Other’ includes errors that do not fall into any other categories, e.g., ‘2 AAs replaced by 4 AAs’.

| Type of sequencing error | pNovo 3 | SMSNet | PointNovo |
| --- | --- | --- | --- |
| Number of total predictions | 16170 | 23227 | 22417 |
| Number of total errors | 14860 | 20554 | 19240 |
| Inversion first 3 AAs | 5.5% | 5.3% | 5.0% |
| Inversion last 3 AAs | 2.4% | 4.4% | 6.7% |
| Inversion first & last 3 AAs | 0.2% | 0.4% | 1.0% |
| 1 AA replaced by 1 AA or 2 AAs | 10% | 5.4% | 5.6% |
| 2 AAs replaced by 2 AAs | 7.0% | 4.5% | 4.2% |
| 3 AAs replaced by 3 AAs | 6.1% | 3.7% | 3.6% |
| 4 AAs replaced by 4 AAs | 2.5% | 3.2% | 2.8% |
| 5 AAs replaced by 5 AAs | 2.6% | 2.6% | 2.1% |
| 6 AAs replaced by 6 AAs | 3.1% | 2.3% | 2.2% |
| More than 6 AAs wrong | 44% | 57% | 55% |
| Other | 14% | 10% | 10% |

**Table 4.** Summary of *de novo* assembly results on light chains of three antibody datasets using *de novo* peptide sequencing tools and the de Bruijn assembler ALPS (k=7). We used the top 20 contigs to compare the length, coverage, and accuracy of mapped contigs. Mapped contigs must be aligned to the reference protein sequence. The longest contig describes the maximum length of all generated contigs. Sequence coverage was calculated as the percentage of amino acids of the complete protein sequence that were covered by at least one contig. Accuracy was calculated as the percentage of all protein sequence calls that were labeled correctly.

|  | IgG1 LC (216 AA) | WlgG1 LC (219 AA) | Herceptin LC (214 AA) |
| --- | --- | --- | --- |
| **pNovo 3** |  |  |  |
| Mapped contigs | 11 | 6 | 6 |
| Longest contig | 28<br>(12.96%) | 23<br>(10.50%) | 51<br>(23.83%) |
| Sequence coverage | 185<br>(85.65%) | 78<br>(35.62%) | 170<br>(79.44%) |
| Sequence accuracy | 154<br>(83.24%) | 75<br>(96.15%) | 154<br>(90.59%) |
| **SMSNet** |  |  |  |
| Mapped contigs | 10 | 5 | 8 |
| Longest contig | 51<br>(23.61%) | 61<br>(27.86%) | 67<br>(31.30%) |
| Sequence coverage | 196<br>(90.74%) | 200<br>(91.32%) | 208<br>(97.20%) |
| Sequence accuracy | 171<br>(87.24%) | 190<br>(95.00%) | 183<br>(87.98%) |
| **PointNovo** |  |  |  |
| Mapped contigs | 7 | 3 | 6 |
| Longest contig | 51<br>(23.61%) | 108<br>(49.32%) | 75<br>(35.05%) |
| Sequence coverage | 205<br>(94.91%) | 204<br>(93.15%) | 212<br>(99.07%) |
| Sequence accuracy | 187<br>(91.22%) | 191<br>(93.63%) | 190<br>(89.62%) |

Further, we evaluated the assembly method on the heavy chains of our evaluated datasets (Supplementary Table S3). Here, we observed a lower sequence coverage and accuracy across all tools. Using PointNovo and ALPS, we achieved a sequence coverage of 64.81% (Herceptin) up to 97.98% (IgG1). We encountered multiple short overlapping contigs, which would make a full-length assembly more difficult without using additional tools. Moreover, these contigs include multiple mismatches, gaps, and were only partly aligned to the target sequence (Supplementary Figure S17). The performance on Herceptin was generally lower since the heavy and light chain were not analyzed separately. Hence, peptides from different chains could be assembled together. Still, on the light chain of Herceptin, we achieved a sequence coverage of 99.07% using PointNovo.

Despite the challenges of full *de novo* protein sequencing, we were able to correctly assemble functionally important subregions, namely the variable region and the CDRs, with the use of PointNovo and ALPS (Table 5). We identified the corresponding CDRs for each antibody using the Natural Antibody database (52). The CDRs were 100% correctly predicted on the light chain of IgG1. The heavy chain was correctly assembled except for a single misidentification on CDR3. The incorrect sequence assignment included mismatches between amino acids with an identical mass (e.g., Q & GA; deamidated N & D; deamidated Q & E).

**Table 5.** Sequence coverage and accuracy of the identified CDRs of three antibodies using PointNovo and ALPS. Sequence coverage was calculated as the number and percentage of amino acids of the CDR that were covered by at least one contig. Accuracy was calculated as the percentage of all CDR sequence calls that were labeled correct. Number in the brackets shows the percentage of correct coverage and accuracy compared to the target sequence. Misidentifications between leucine and isoleucine were not counted as incorrect. CDRs were identified via the Natural Antibody database (research.naturalantibody.com/pad).

| Antibody | CDR1 |  | CDR2 |  | CDR3 |  |
| --- | --- | --- | --- | --- | --- | --- |
|  | Cov. | Acc. | Cov. | Acc. | Cov. | Acc. |
| IgG-1 Human HC | 10 of 10<br>(100%) | 10 of 10<br>(100%) | 7 of 7<br>(100%) | 7 of 7<br>(100%) | 9 of 9<br>(100%) | 8 of 9<br>(88.8%) |
| IgG-1 Human LC | 6 of 6<br>(100%) | 6 of 6<br>(100%) | 3 of 3<br>(100%) | 3 of 3<br>(100%) | 9 of 9<br>(100%) | 9 of 9<br>(100%) |
| WlgG Mouse HC | 0 of 8<br>(0%) | 0 of 0<br>(0%) | 7 of 7<br>(100%) | 6 of 7<br>(85.7%) | 12 of 12<br>(100%) | 11 of 12<br>(91.7%) |
| WlgG Mouse LC | 11 of 11<br>(100%) | 10 of 11<br>(90.9%) | 3 of 3<br>(100%) | 3 of 3<br>(100%) | 9 of 9<br>(100%) | 9 of 9<br>(100%) |
| Herceptin HC | 8 of 8<br>(100%) | 8 of 8<br>(100%) | 8 of 8<br>(100%) | 7 of 8<br>(100%) | 9 of 13<br>(69.2%) | 2 of 9<br>(22.2%) |
| Herceptin LC | 6 of 6<br>(100%) | 5 of 6<br>(83.3%) | 3 of 3<br>(100%) | 3 of 3<br>(100%) | 9 of 9<br>(100%) | 8 of 9<br>(88.8%) |

On the heavy chain of IgG1, we achieved a coverage of and entirely correct predictions of CDR1 and CDR2. At CDR3, we observed a mismatch of T instead of F (Figure 8). We were able to cover the complete sequence using only three contigs, apart from 6 AAs in FR1 and 2 AAs in FR4. On Herceptin, we achieved a lower coverage and accuracy, especially on CDR3 of the heavy chain due to the combined analysis of the heavy and light chain. The results on the variable regions and CDRs highlight the potential of deep learning-based *de novo* sequencing to identify unique antibody sequences.

**Figure 8.**
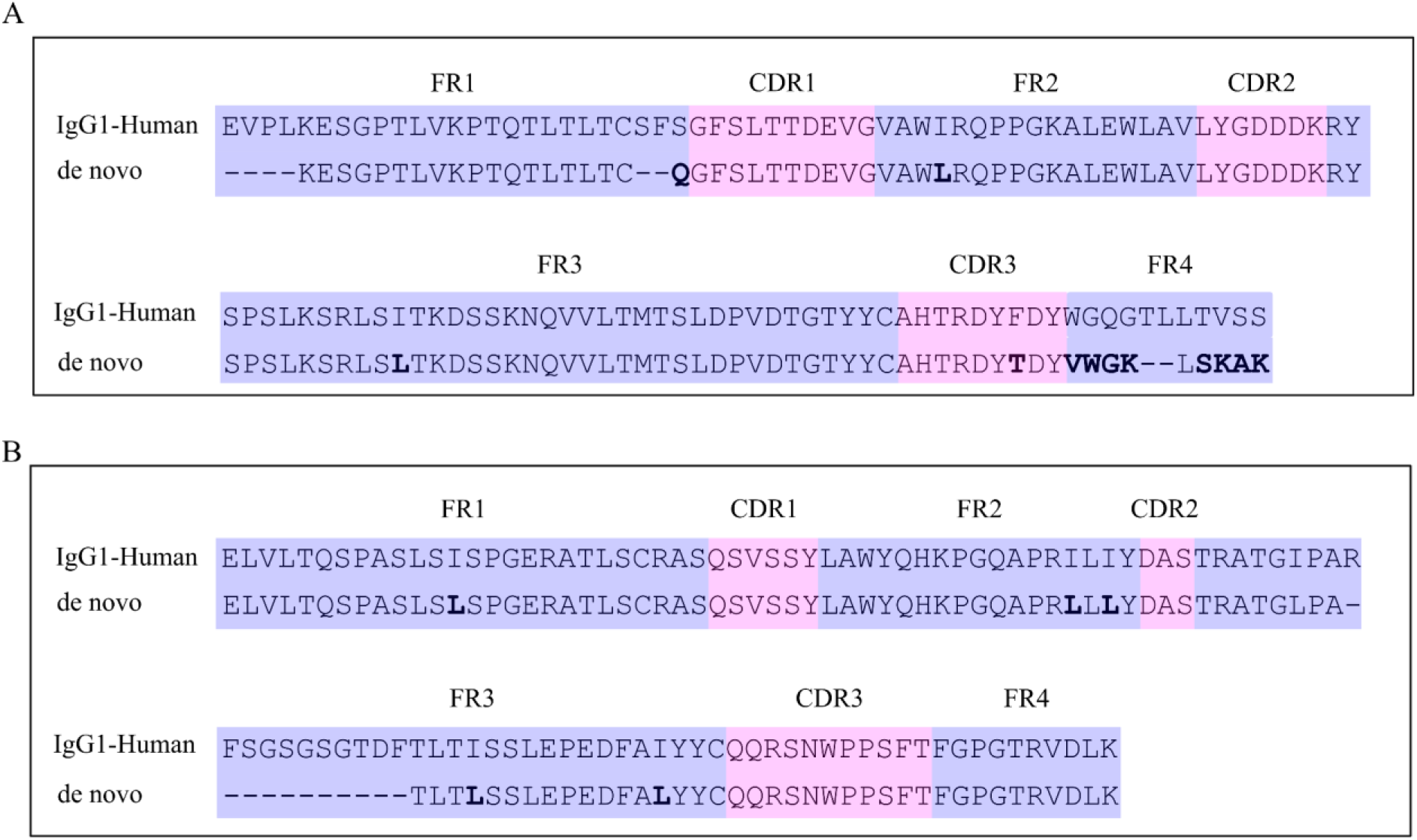
*De novo* assembly of the variable region of the heavy chain (A) and light chain (B) of IgG1-Human. We used all available enzymes for *de novo* sequencing with PointNovo and assembly with ALPS. We aligned the resulting contigs (5 for Heavy chain and 4 for Light Chain) against the target protein to reconstruct the antibody sequence. CDRs are marked in pink, while framework regions (FRs) are marked in blue. Mismatches are indicated by bold characters. Gaps are indicated by dashes. FRs and CDRs were numbered according to the Natural Antibody database (research.naturalantibody.com/pad).

In summary, the biggest issue lies in short contigs, which alone only cover subregions of the whole sequence leading to incomplete assemblies, particularly on long protein sequences such as the heavy chains of our evaluated antibodies. While recently developed deep learning tools improved the accuracy on peptide level and showed an increased sequence coverage in comparison to spectrum-graph-based tools, *de novo* protein sequencing remains a demanding task. Thus, additional tools and assembly strategies for protein identification are necessary to correctly align short contigs and improve the assembled results from *de novo* peptide sequencing.

## 4 Discussion

In this study, we reviewed *de novo* sequencing algorithms and applied them to the assembly of monoclonal antibodies. We compared the performance of five recently developed and commonly used *de novo* peptide sequencing tools, namely Novor, pNovo 3, DeepNovo, SMSNet, and PointNovo.

Statistical analysis on amino acid and peptide levels revealed that the recently developed tools SMSNet and PointNovo achieved a high peptide recall on different enzymatic datasets (Figure 2, Figure 3, Supplementary Figure S3). Similar to previous observations (21,53), deep learning-based algorithms, which employ encoder-decoder networks, predict a higher amount of correct peptide sequences compared to conventional spectrum-graph-based methods. Beside the use of the encoder-decoder architecture, the number of fragment ions for predicting each amino acid residue play another important role. While Novor and DeepNovo use 8 ion types for predicting each position (y, y(2+), y-NH2, y-H2O, b, b(2+), b-NH2, b-H2O), SMSNet and pNovo 3 take 9 ion types into consideration for inferring peptides from spectrum peaks. Moreover, PointNovo examines 12 ion types to calculate theoretical m/z values at each prediction step. Checking additional ion types can help interpreting spectra, where b- and y-ions are missing (Figure 6, Supplementary Figure S18). Another crucial factor for retrieving the correct peptide sequence is the resolution of the mass instrument and consequently the ability of *de novo* sequencing tools to make use of such high-resolution spectra. Ambiguities between amino acids with similar mass (e.g. Q & K; oxidized M & F; AG & Q) cannot be resolved correctly on mass spectra with a fragment ion error tolerance of 0.1 Da and make MS/MS of higher resolution necessary (54). As Qiao *et al*. pointed out, spectra of higher resolution led to a higher computational complexity for analyzing MS/MS data with *de novo* sequencing algorithms. DeepNovo and SMSNet would need to discretize spectra with a higher resolution parameter, which would increase the computation and memory demand, whereas PointNovo can handle high-resolution spectra without increasing its computational complexity (21). However, particular amino acids cannot be resolved even with spectra and tools with higher resolution (e.g., I & L; Q & AG; deamidated N & D). Here, additional methods are necessary to retrieve the correct amino acid sequence. Discrimination of the isomeric residues isoleucine and leucine cannot be achieved via tandem mass spectrometry but require MS3 fragmentation (55,56).

The performance of pNovo 3 demonstrated that traditional *de novo* sequencing, coupled with a learning-to-rank framework and deep learning architecture from pDeep, still achieves a higher precision than DL-based approaches. pNovo 3 was able to achieve a remarkably higher amino acid precision across all enzymes (Figure 2) and generate less sequence errors than SMSNet and PointNovo (Table 2). Furthermore, the approach from pNovo 3 achieved the highest peptide recall on asp-N and chymotrypsin of IgG1-Human-HC (Supplementary Figure S3), due to its enzyme-specific pre-trained models. Cross-enzyme performance is a crucial quality of *de novo* sequencing methods in bottom-up proteomics. Yet, most publications regarding *de novo* sequencing tools rarely address the predictive abilities of non-tryptic proteolytic enzymes and focus on tryptic datasets due to their availability and well-established usage. Qiao *et al*. observed that enzyme-specific models had a notable influence on the performance and recommended training a separate model for each enzyme (21). However, training different models for over 6 enzymes can be a demanding task, especially if the data is only partly available or coming from various sources with unequal experimental setups. Lee *et al*. applied DeepNovo for metaproteomic data analysis by training a model on 5 million peptide-spectrum matches from multiple bacteria (57). In Supplementary Figure 1 of their publication, Lee *et al*. observed the increased peptide recall when using a higher amount of MS/MS spectra for the training procedure. Although our training data included mainly tryptic peptides, a relatively high number of non-tryptic peptides were identified by PointNovo, SMSNet, and DeepNovo (Figure 2). Karunratanakul *et al*. made a similar observation, where SMSNet was able to discover a large number of non-tryptic HLA-antigens, while 95% of its training data consisted of tryptic peptides (20). We conclude that DL tools can still be applied to different enzymatic datasets, although the performance will vary based on the cleavage pattern of the trained dataset. The deployment of a higher number of enzyme-specific datasets and models would benefit the field of *de novo* sequencing in bottom-up proteomics.

Previous evaluations of *de novo* sequencing tools have observed an increased accuracy of these algorithms on simulated MS/MS spectra compared to real datasets (12,29), suggesting that the bottleneck for *de novo* peptide identification lies in the quality of the provided data. As shown in our analysis, all evaluated tools show a higher peptide recall on spectra with less missing fragment ions (Figure 5). McDonnell *et al*. (2022) pointed out that over 72% of all evaluated peptide spectrum matches lack at least one fragment ion. We observed an even higher value of 90.51% of all spectra lacking at least one fragment ion. While newly developed tools demonstrate the potential of *de novo* sequencing, advanced post-processing steps are necessary to improve their accuracy. The DL-based tools SMSNet and PointNovo generated a higher number of completely incorrect peptides in comparison to pNovo 3 (Table 2). As Yang et al. pointed out, DL models are directly learned from the MS/MS data and do not rely on well-designed features, which could help reduce the error frequency. Furthermore, the authors reported that even DL-based approaches have difficulties in distinguishing similar peptides with long gapped subsequences concluding that the quality of MS/MS data is a bottleneck of successful peptide prediction (33). Current state-of-the-art tools include hybrid approaches to address the issue of incomplete fragmentation ions in mass spectra (20,21). SMSNet includes a sequence-mask-search, which replaces segments of low confidence amino acids with their total residue mass, using database search to identify the missing subsequence. PointNovo incorporates a database search module using Percolator (58). In addition, it is worth mentioning that several methods were published, discussing how to improve the encoder-decoder paradigm of deep learning tools in proteomics (53,59). Fei pointed out that deep neural networks face difficulties on tandem mass spectra with incomplete fragment patterns. Multiple authors have confirmed that a considerable amount of *de novo* sequencing errors occurs at the N-terminal ends due to the absence and low intensity of fragment ions (60,61). Hence, Fei developed a retrieve-and-revise framework to compensate for low-quality spectra. His peptide identification model, which relies on a reference database, was able to outperform current state-of-the-art algorithms (53). In another study, Fei pointed out that the deep neural network approach was limited by generating full predictions one by one left to right. The author developed a tag-based method named DeepTag, which first predicts high-confidence tags and subsequently generates the whole peptide sequence by prolongation of the identified tags (62). Ge *et al*. proposed the use of deep residual shrinkage networks for their *de novo* sequencing method DePS to improve the accuracy on noisy spectra with missing fragmentation ions. Their implementation improved the extraction of features from MS/MS spectra and outperformed DeepNovoV2 on multiple datasets (63).

Another promising approach to boost the number of correct predictions lies in complementary methods using predictions of different *de novo* sequencing tools. Combining the predictions of SMSNet and PointNovo resulted in a higher number of correct identifications (Figure 4). Currently, only few post-processing approaches have been described for deep learning predictions in mass spectrometry. Li *et al*. developed DeepRescore, a post-processing tool for database search, which uses retention time and simulated MS/MS spectra to rerank peptide-spectrum matches. Using public datasets, the authors demonstrated how their rescoring method increases sensitivity and reliability compared to state-of-the-art methods (64). A similar approach would be beneficial for improving the quality of *de novo* sequencing based on recently developed DL tools. To our knowledge, PostNovo is currently the only post-processing tool for *de novo* sequencing results from different algorithms. Using the predictions from DeepNovo, Novor, and PepNovo, PostNovo can improve the accuracy by re-scoring and re-ranking steps. Its sequence-recall exceeds the best-performing tools at that time, DeepNovo and Novor, by a factor of 7 to 15 at an FDR of 10% (61). Furthermore, using information of missing fragmentation gaps such as in pNovo 3 can help to reduce the number of incorrect predictions for long peptide sequences (33). Here, the missing fragmentation sites are designed as features to calculate the probability of losing fragment ions between different combinations of amino acids and use this information in a learning-to-rank model. Thus, DeepRescore (64), MS2Rescore (65) and pNovo 3 (33) demonstrate the advantages of DL-based spectrum prediction tools for a re-ranking approach of PSMs. The prediction of mass spectra and retention time, gap features, and the consensus results from multiple tools contain useful information to develop a scoring metric for accurate *de novo* sequencing.

In addition to the algorithmic solutions described above, the accuracy of *de novo* sequencing can be increased by alternative experimental setups, including the use of complementary fragmentation techniques or multiple proteolytic enzymes. For instance, using alternative fragmentation methods, such as HCD, CID, and ETD, is a reliable method to increase the correct detection of peptides (10,22), for which algorithms like pNovo+ (66) and CompNovo (67) have been developed. While the use of multiple proteases with different cleavage sites leads to a higher protein coverage due to overlapping peptide fragments (10,23,68), it comes along with severely prolonged sample preparation and an increased analysis time. Digestion with cleavage-specific enzymes such as trypsin can result in numerous peptides that are not suited for MS-based analysis due to their length, hydrophobicity, or (poor) ionization. This can limit the chance of achieving a protein coverage of 100% (25). Moreover, microwave-assisted acid hydrolysis (24,50), the complementary use of the mirror proteases trypsin and Ac-LysargiNase (33), the application of trypsin and ProAlanase (69), and the complementary use of lys-C and lys-N (51) have been described as alternative experimental setups for bottom-up proteomics. Here again, multiple tools have been developed for alternative proteases, e.g., Lys-Sequencer (51) for lys-C and lys-N data, pTa (24) and MuCS (25) for microwave-assisted acid hydrolysis.

Despite the ongoing effort and progress in *de novo* peptide sequencing, reliable protein assembly is still a demanding task. Our findings show that the ability of database-independent approaches of full-length protein assembly is limited even when using multiple contigs and different *de novo* sequencing tools (Table 4). The longest generated contig only covers at best 21.82% of the heavy chain and 49.32% of the light chain. Using PointNovo and ALPS, we accomplished a sequence coverage of 93.15% to 99.07% on the light chains of our evaluated antibodies. However, additional database tools or homology algorithms are necessary to correctly assemble multiple short contigs to long protein sequences. The software package SPIDER uses database search and *de novo* sequencing tags to identify homologs of the target protein by minimizing the sequencing errors and potential number of mutations (28). BICEPS follows a similar approach by using Bayesian statistical criteria to identify the optimal trade-off between the database and spectral information to identify proteins (70). Hence, homology search can be combined together with *de novo* sequencing to improve the discovery of protein sequence information and overcome problems caused by mass segment errors (22,71,72). Commercial software packages like PEAKS AB (23) and Supernovo (9) use antibody germline sequences as a starting point together with *de novo* sequencing results to identify monoclonal antibodies. Supernovo employs *de novo* peptide sequencing, database search, *in silico* genetic recombination, and a final sequence assembly for an automatic antibody sequence prediction. As Olsen *et al*. observed, 40% of all antibodies in the Observed Antibody Space (OAS) database (73) are missing the first 15 amino acids positions. In addition to the before-mentioned homology tools, antibody-specific language models, such as AbLang, can help to restore missing residues of full protein sequences caused by sequencing errors without using a germline template sequence (74). Thus, the development of publicly available frameworks and pipelines for automated assembly of *de novo* peptide sequencing results from recently developed algorithms would improve the reliable usability of *de novo* sequencing for full antibody assembly. In this context, it is also questionable whether database-independent sequencing of whole antibody sequences is necessary. In case there is access to the DNA of the hybridoma cell, the cDNA of the variable domain is usually sequenced and the antibody is then recombinantly produced with a Fc domain of the researcher’s choice (75). Moreover, a large part of antibody sequences does not show a high variability and can be retrieved with database and homology search (9,23,76). Therefore, obtaining the whole antibody sequence on the protein level by *de novo* sequencing may be an ambitious goal that is not even needed in many cases. Reducing the complexity by only sequencing the variable domains on the protein level and employing effective homology methods can already be sufficient for retrieving complete antibody sequences.

## 5 Conclusion

Recent advances in machine learning, the availability of large datasets, and high-resolution mass spectrometry have contributed to the overall progress of *de novo* sequencing of antibody sequences. We observed that deep learning-based algorithms, specifically SMSNet and PointNovo, show an increased number of correct peptide predictions compared to spectrum-graph-based approaches. A complementary approach using multiple *de novo* sequencing tools would further increase the accuracy of database-independent peptide identification. However, long peptide sequences and missing fragment ions still pose a challenging task to all *de novo* sequencing tools. Besides algorithmic optimization, the choice of alternative experimental setups and proteolytic enzymes would help to reduce the number of noise peaks, missing fragment ions, and peptide length. The full assembly of *de novo* peptide sequences without the use of any additional database or homology search tool remains a challenge on long antibody sequences. While we were able to identify certain subregions of our evaluated antibodies, retrieving the complete antibody sequence without additional tools could not be resolved sufficiently. Our findings point to the need of additional frameworks and pipelines for reconstructing and assembling full antibody sequences. The development of publically available hybrid approaches using *de novo* sequencing and homology search would benefit the identification and assembly of full-protein sequences.

## Key Points

- A comprehensive review of *de novo* sequencing tools in proteomics is provided that aim to solve the challenge of antibody sequencing and subsequent assembly.
- An improved sensitivity of deep learning-based tools was found in comparison to classical *de novo* sequencing algorithms such as spectrum graph-based algorithms, across various enzymatic datasets of antibodies.
- The number of missing fragmentation sites, noisy spectra, and long peptide sequences pose a limit for all *de novo* sequencing tools
- Database-independent assembly of light chains can be achieved up to a sequence coverage of 99.07% by using the de Bruijn assembler ALPS together with *de novo* peptide predictions from PointNovo.
- Further development of freely available and automatized pipelines for an accurate assembly of peptide predictions is necessary to successfully retrieve full antibody sequences.

## Acknowledgement

The authors would like to thank Natthanan Ruengchaijatuporn (Chulalongkorn University) for his help for the re-training process of SMSNet.

## Author contributions

D.B. collected the data and performed the main analysis. T.M. and B.R. supervised the project. D.B. and T.M. wrote the manuscript with the help from B.R., G.T., and M.W. All authors reviewed and approved the manuscript.

## Data availability

Result files and Python code to reproduce the results in this study are available at Figshare (doi.org/10.6084/m9.figshare.20332692).

## Funding

**Denis Beslic** is a research assistant at the Robert Koch Institute in Berlin, Germany. His research interests include machine learning, computational biology, and proteomics data analysis.

**Thilo Muth** is the head of section S.3 (eScience) at the Federal Institute for Materials Research and Testing (BAM) in Berlin, Germany. His research interests include developing bioinformatic methods for protein analytics and providing data-driven applications for material science and engineering.

**Bernhard Y. Renard** is professor for data analytics and computational statistics at Hasso - Plattner-Institute and the University of Potsdam in Potsdam, Germany. His research interests include omics data analysis and explainable machine learning.

**Georg Tscheuschner** is a doctoral candidate at the Federal Institute for Materials Research and Testing (BAM) in Berlin, Germany. His research interests include developing methods for the identification and traceability of monoclonal antibodies and the use of plant viruses as nanotechnological platforms.

**Michael G. Weller** is the head of division 1.5 (Protein Analysis) at the Federal Institute for Materials Research and Testing (BAM) in Berlin, Germany. His research interests include bioanalytical subjects such as metrology of protein quantification, immunoassays, biosensors, microarrays, affinity chromatography, monolithic materials, antibody development, bioconjugation, lab-on-a-chip, and combinatorial peptide libraries. Furthermore, he is active in the design and quality control of bioreagents to overcome the replication crisis.

