## Supplementary Data for "Current state, existing challenges, and promising progress for *de novo* sequencing and assembly of monoclonal antibodies"

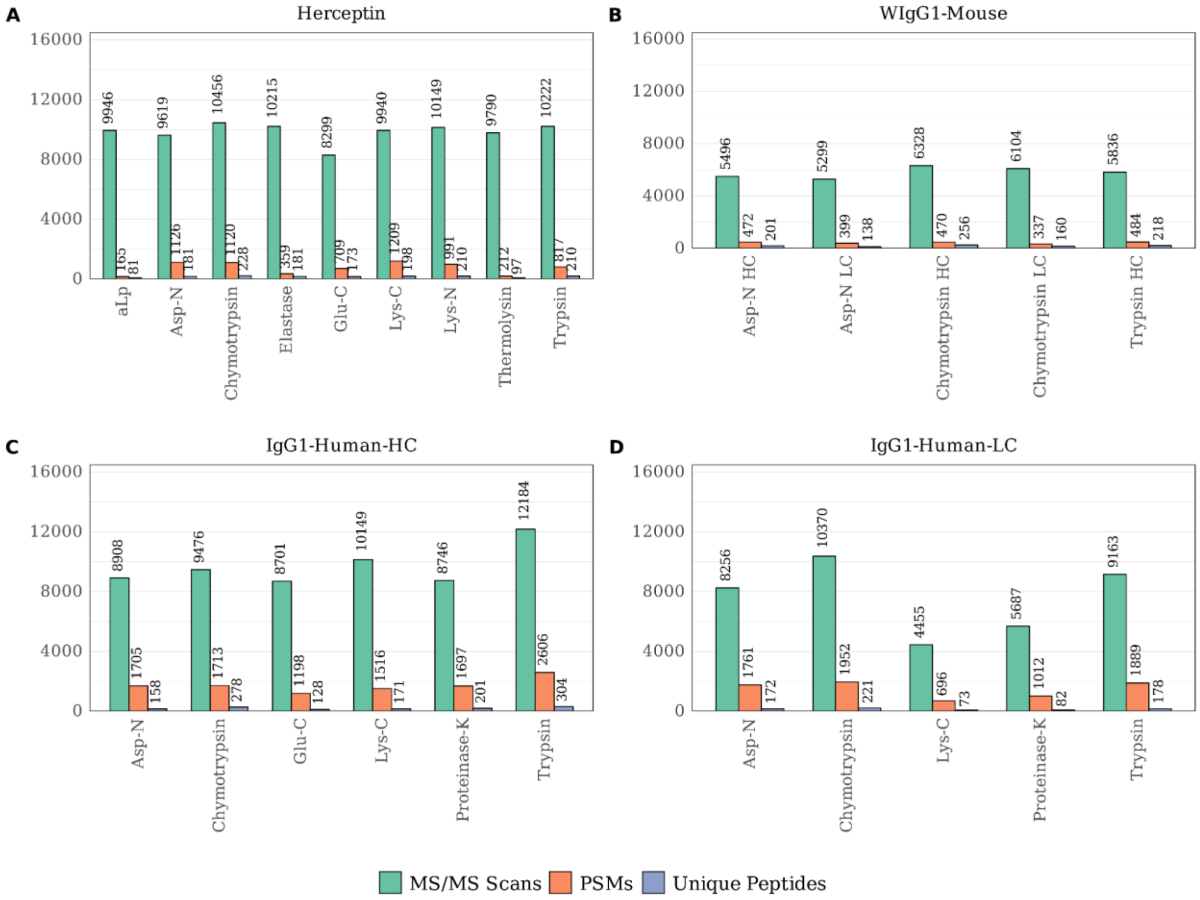


**Supplementary Figure S1** | Scan statistics for each protease digest of Herceptin (A), WIgG1-Mouse (B), the heavy chain (C), and the light chain (D) of IgG1-Human. For each dataset, the number of MS/MS scans (green), the number of peptide-spectrum matches (orange), the number of unique peptides (blue), are displayed.

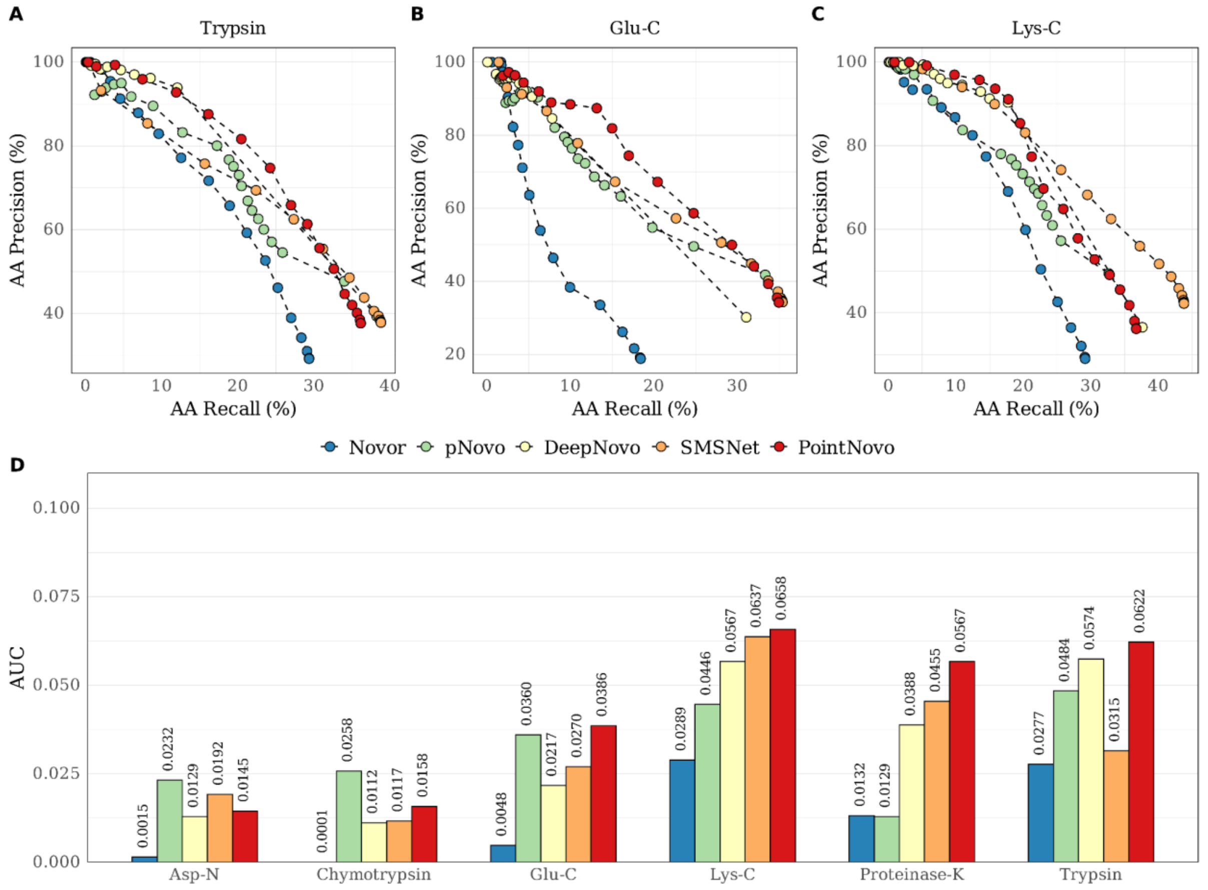


**Supplementary Figure S2** | The precision-recall (PR) curves of Novor, pNovo 3, DeepNovo, SMSNet, PointNovo for trypsin (A), glu-C (B), and lys-C (C) of the IgG1-Human-HC dataset. The area under curve (AUC) of the five algorithms for each PR curve and each enzyme of IgG1-Human-HC (D).

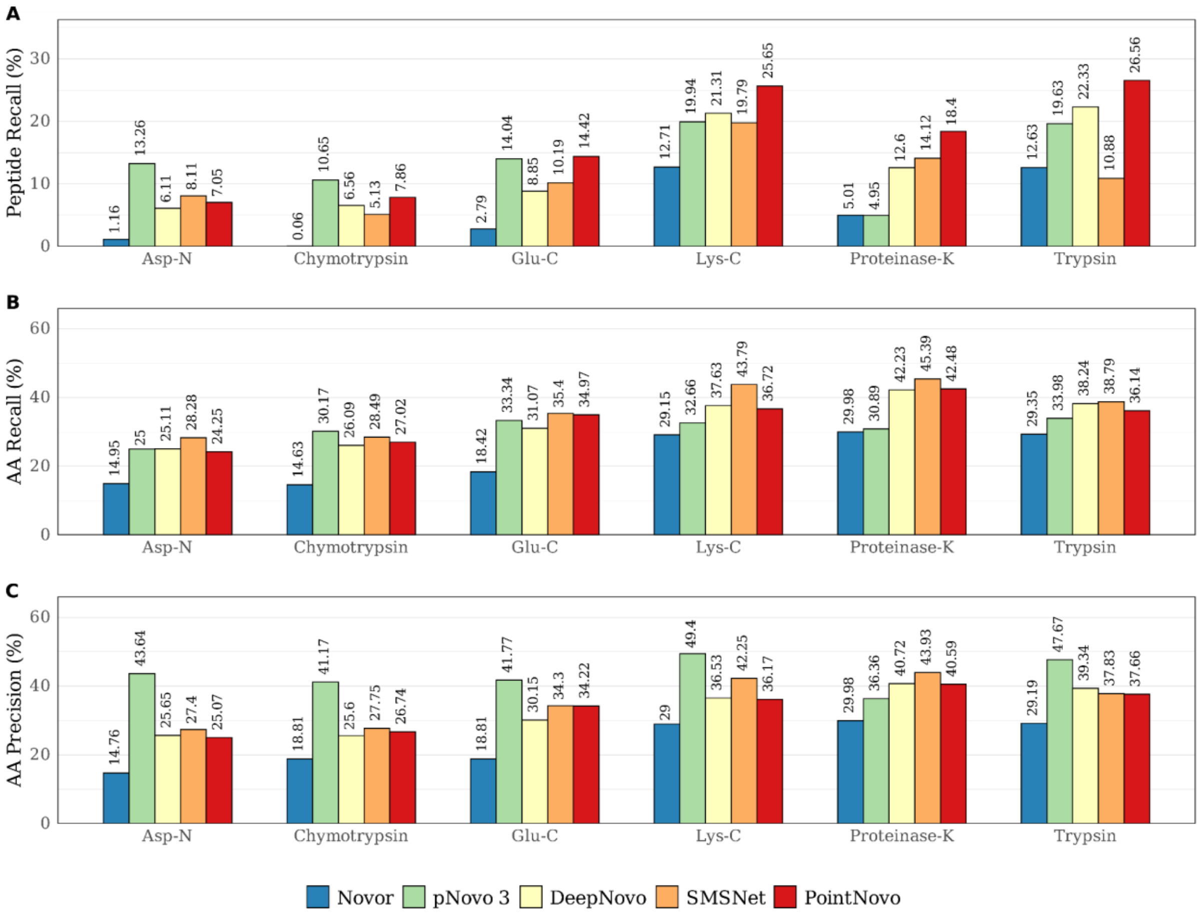


**Supplementary Figure S3** | Total recall and precision of Novor, pNovo 3, DeepNovo, SMSNet and PointNovo across different enzymes on IgG1-Human-HC. (A) Recall at peptide level. (B) Recall at amino acid level. (C) Precision at amino acid level.

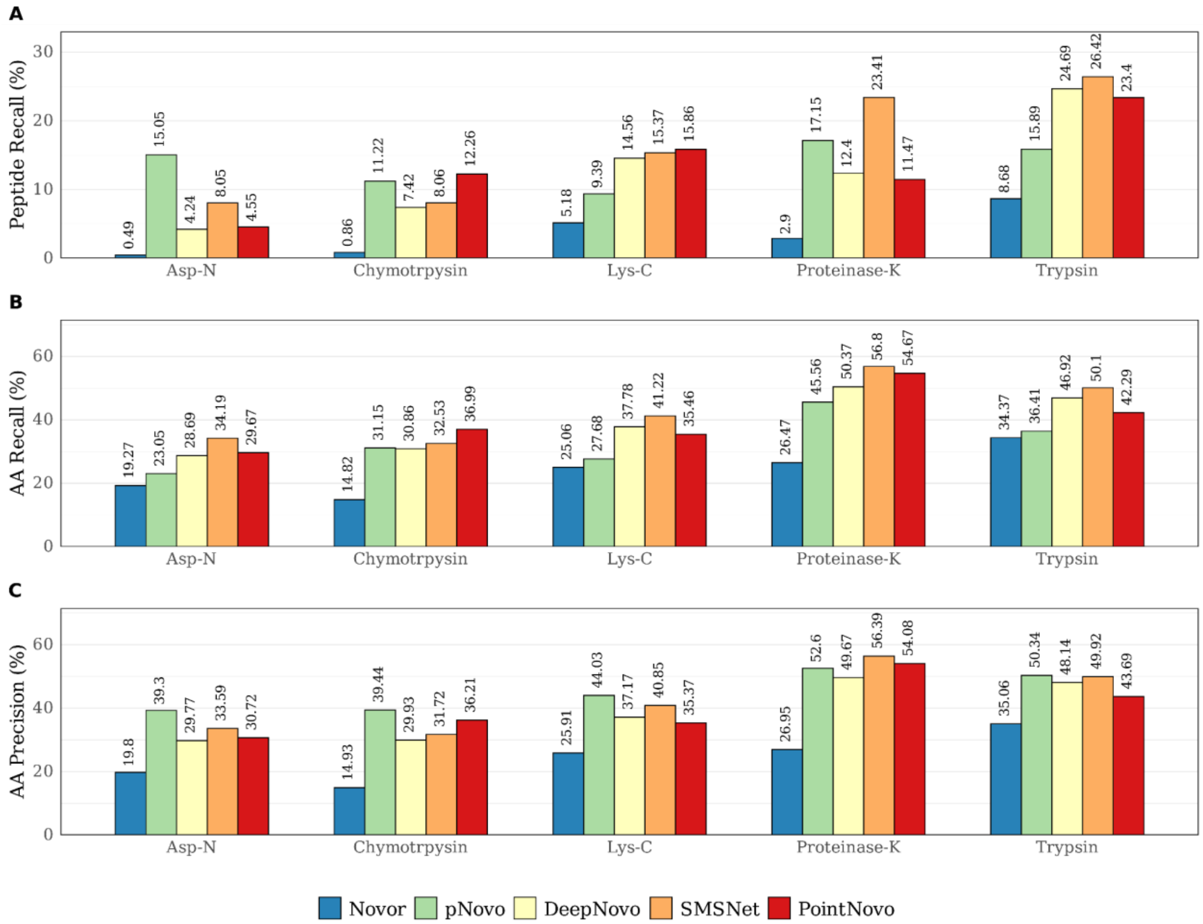


**Supplementary Figure S4** | Total recall and precision of Novor, pNovo 3, DeepNovo, SMSNet and PointNovo across different enzymes on IgG1-Human-LC. (A) Recall at peptide level. (B) Recall at amino acid level. (C) Precision at amino acid level.

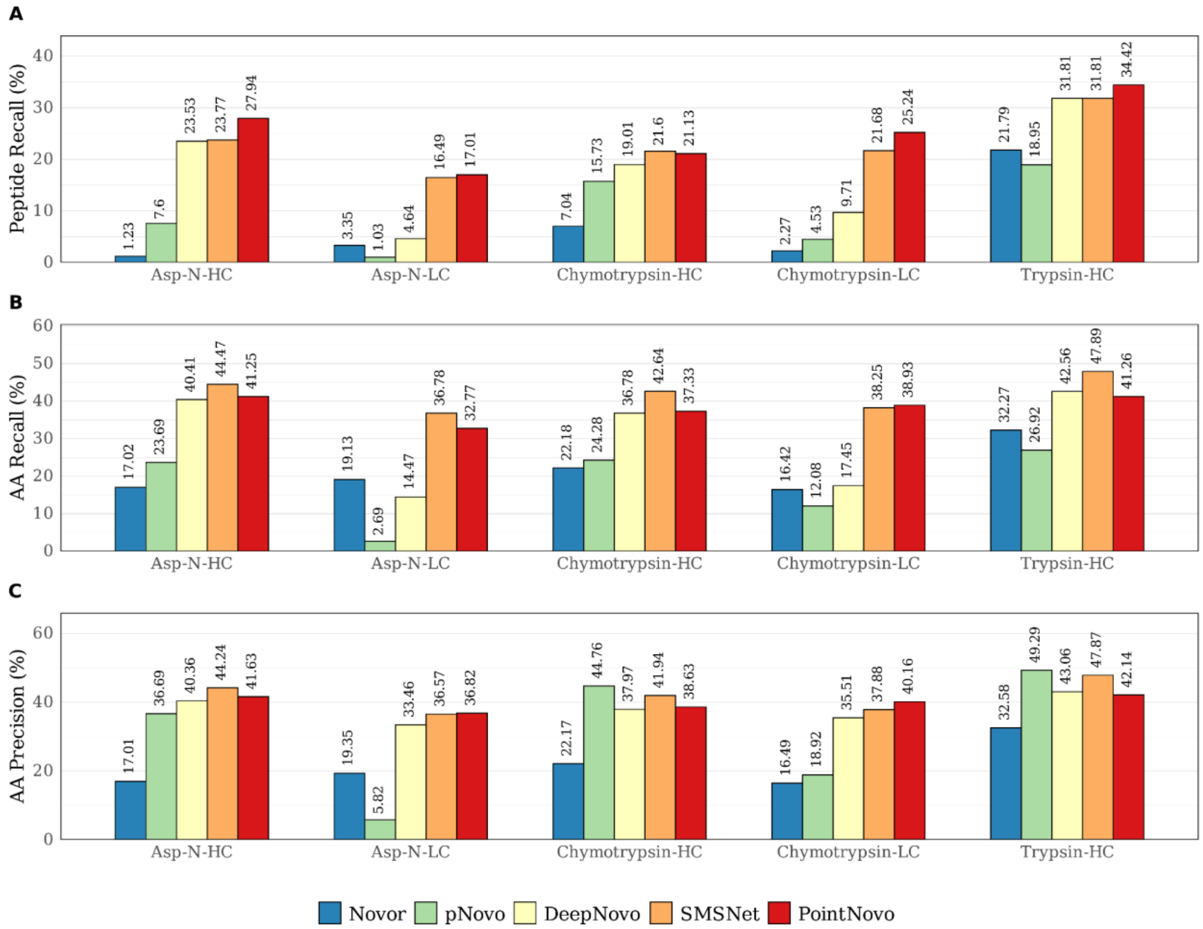


**Supplementary Figure S5** | Total recall and precision of Novor, pNovo 3, DeepNovo, SMSNet and PointNovo across different enzymes on WIgG1-Mouse. (A) Recall at peptide level. (B) Recall at amino acid level. (C) Precision at amino acid level.

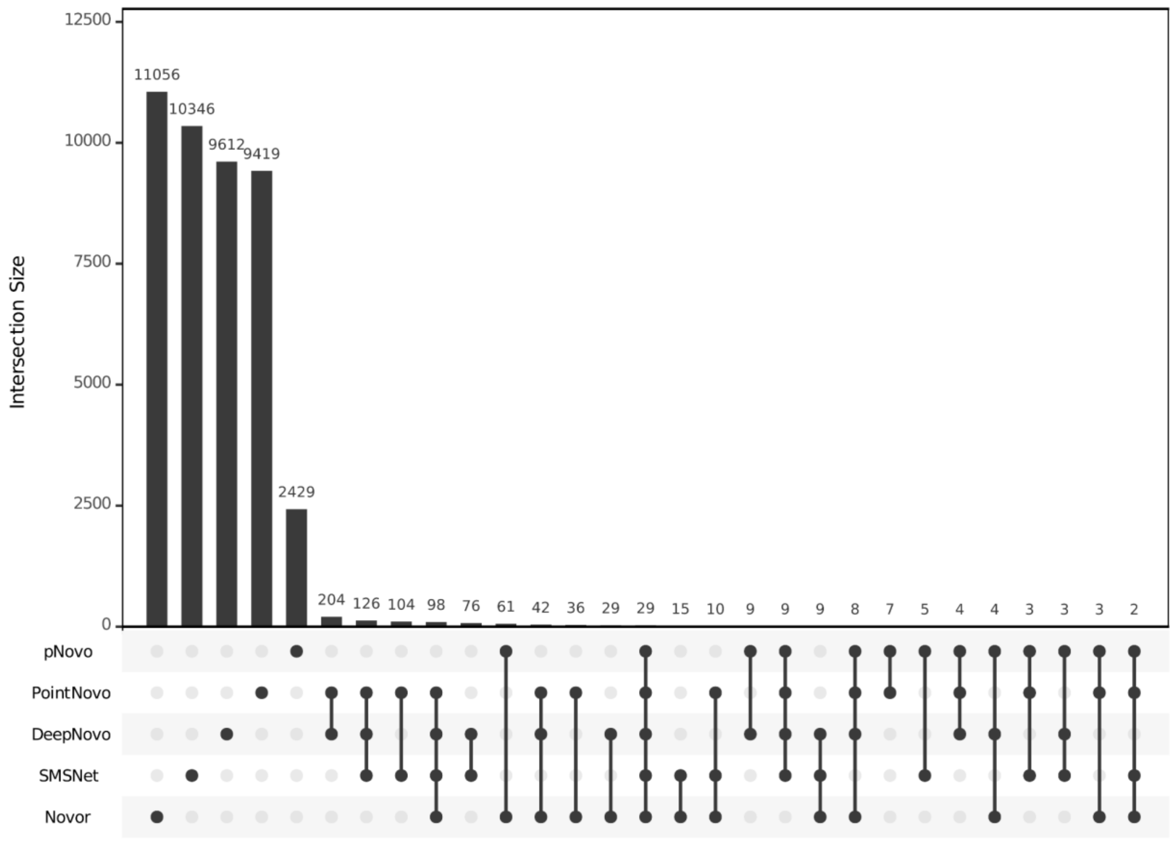


**Supplementary Figure S6** | Upset plot showing the number of commonly identified peptide sequences between different sets of *de novo* sequencing tools for the IgG1-Human-HC Trypsin dataset.

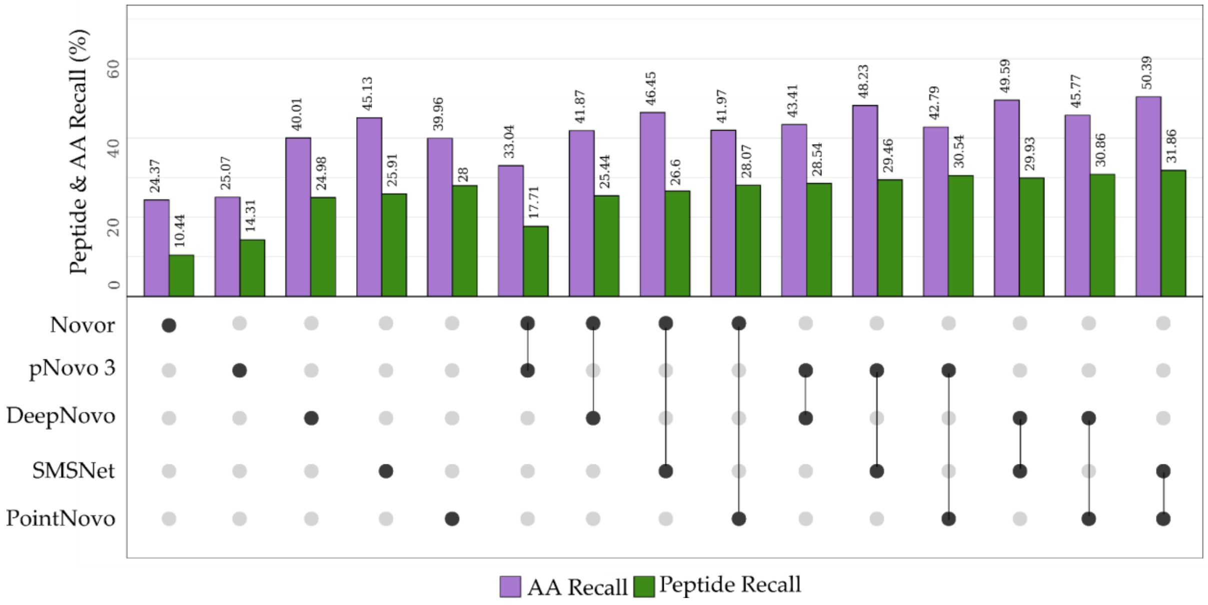


**Supplementary Figure S7** | The AA recall (violet) and peptide recall (green) by each pair of combined *de novo* sequencing results for the WIgG1-Mouse-HC dataset.

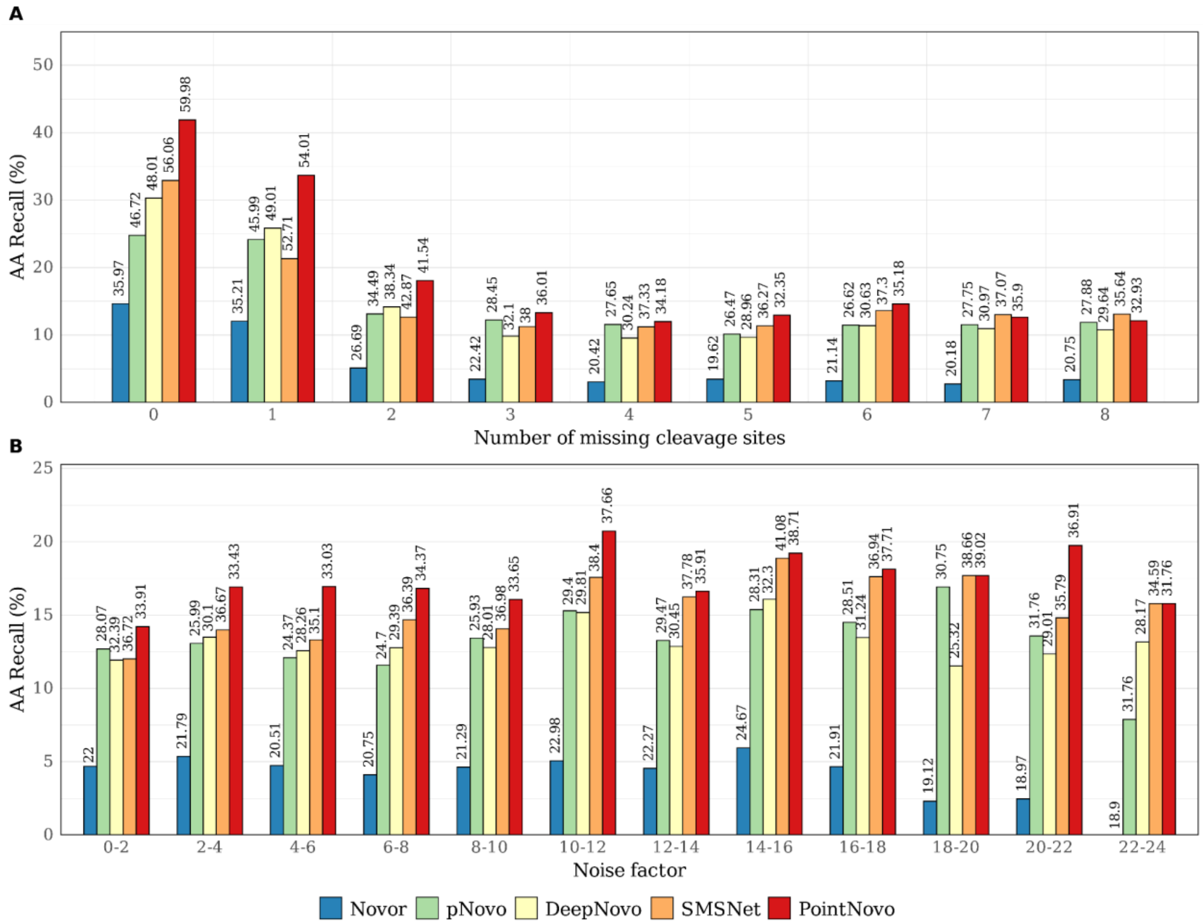


**Supplementary Figure S8** | Total amino acid recall of Novor, pNovo 3, DeepNovo, SMSNet and PointNovo across all datasets for different number of cleavage sites missing (A) and different noise factors (B) of the specific spectra.

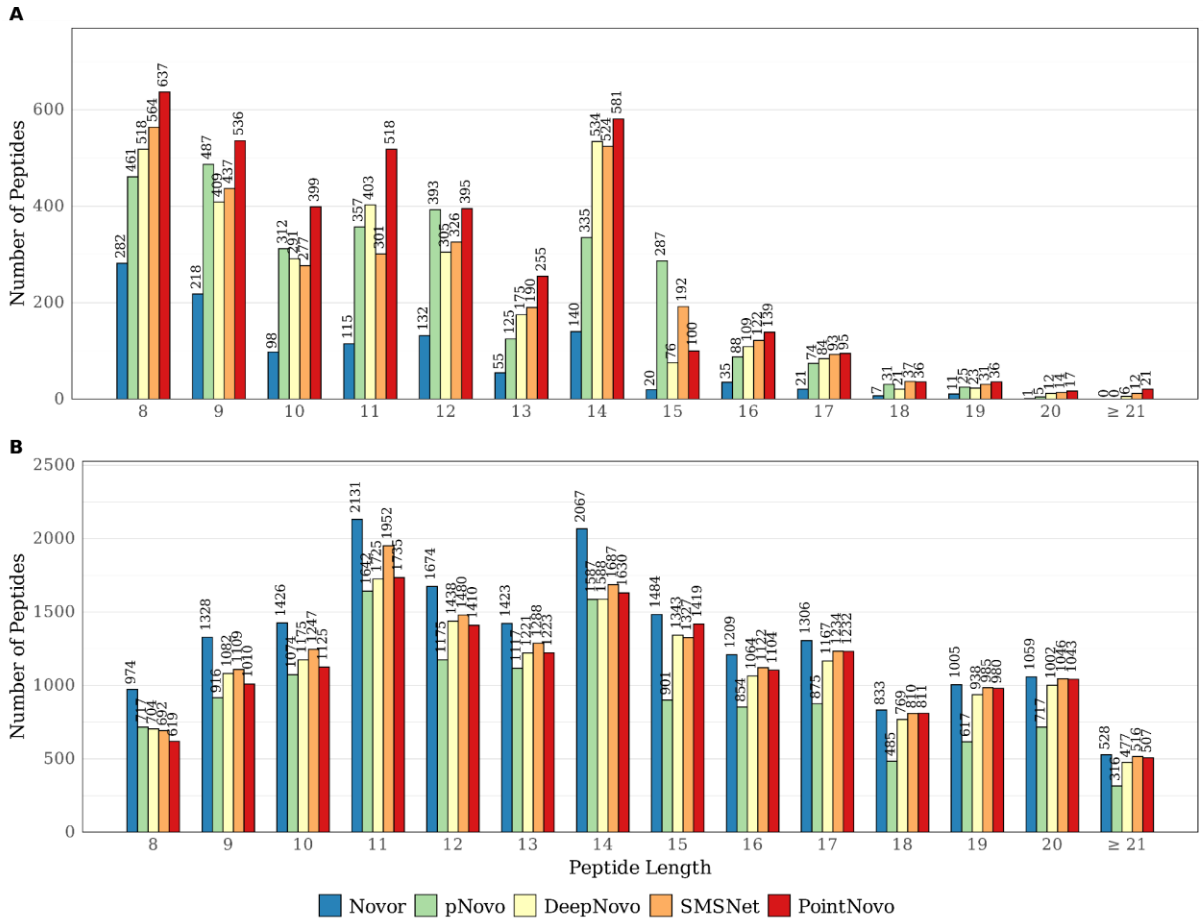


**Supplementary Figure S9** | Peptide length distribution of correct (A) and incorrect (B) predictions of Novor, pNovo 3, DeepNovo, SMSNet and PointNovo on all datasets.

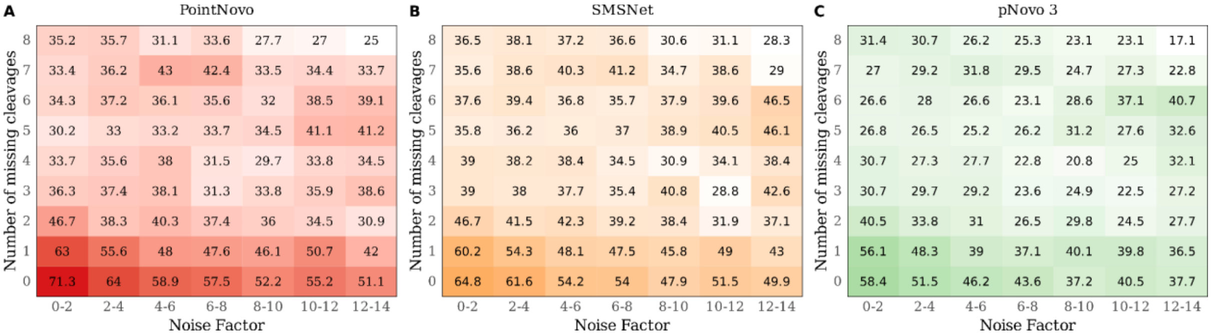


**Supplementary Figure S10** | Heat plot showing amino acid recall for different number of missing cleavages (Y-Axis) and noise factors (X-Axis). Higher AA recall is shown in red for PointNovo (A), orange for SMSNet (B), and green for pNovo 3 (C). Lower AA recall is displayed in white.

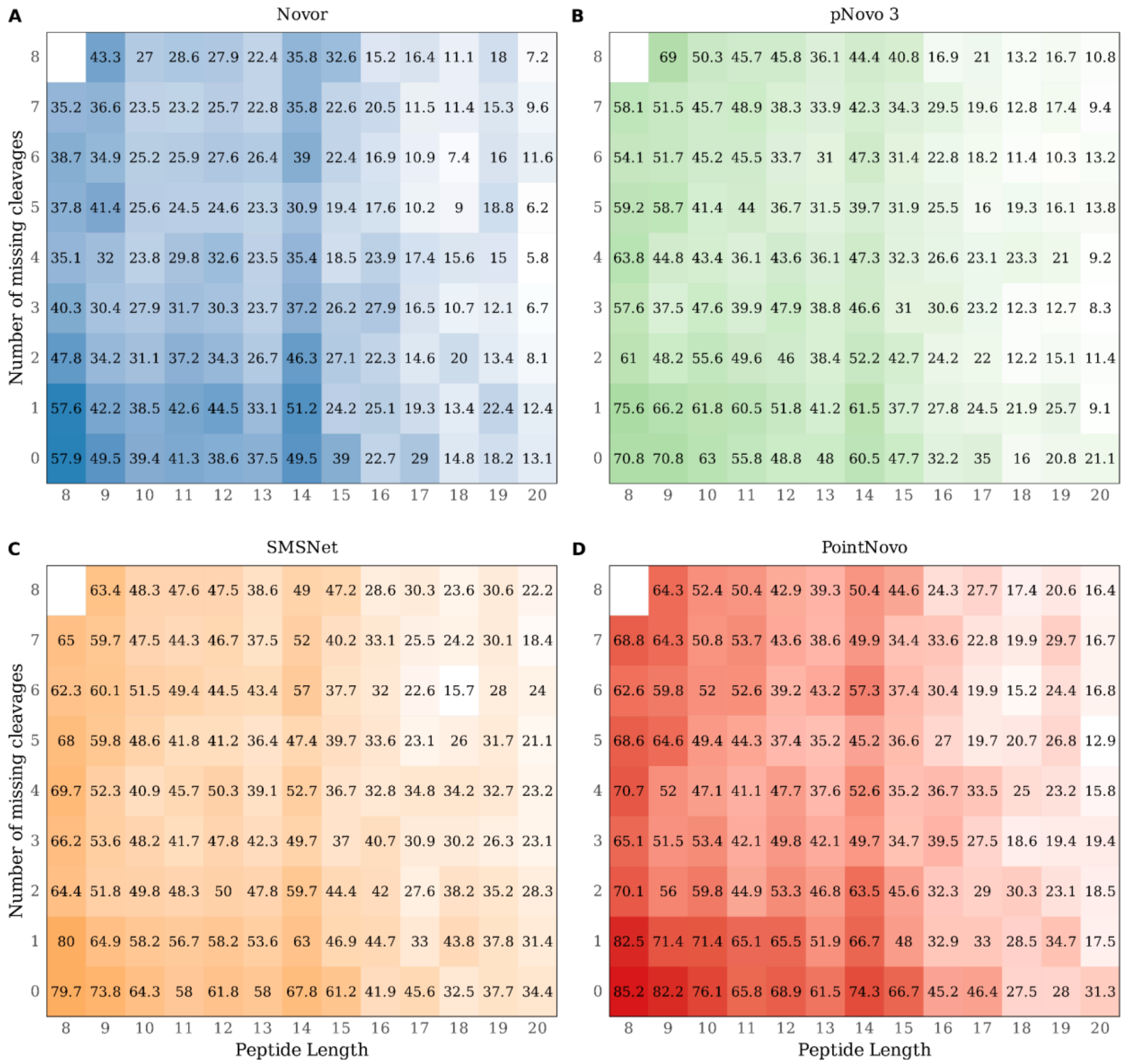


**Supplementary Figure S11** | Heatmap showing amino acid recall for the number of missing cleavages (Y-Axis) and peptide lengths (X-Axis) across all datasets. Higher AA recall is shown in blue for Novor (A), green for pNovo 3 (B), orange for SMSNet (C), and red for PointNovo (D). Lower AA recall is displayed in white.

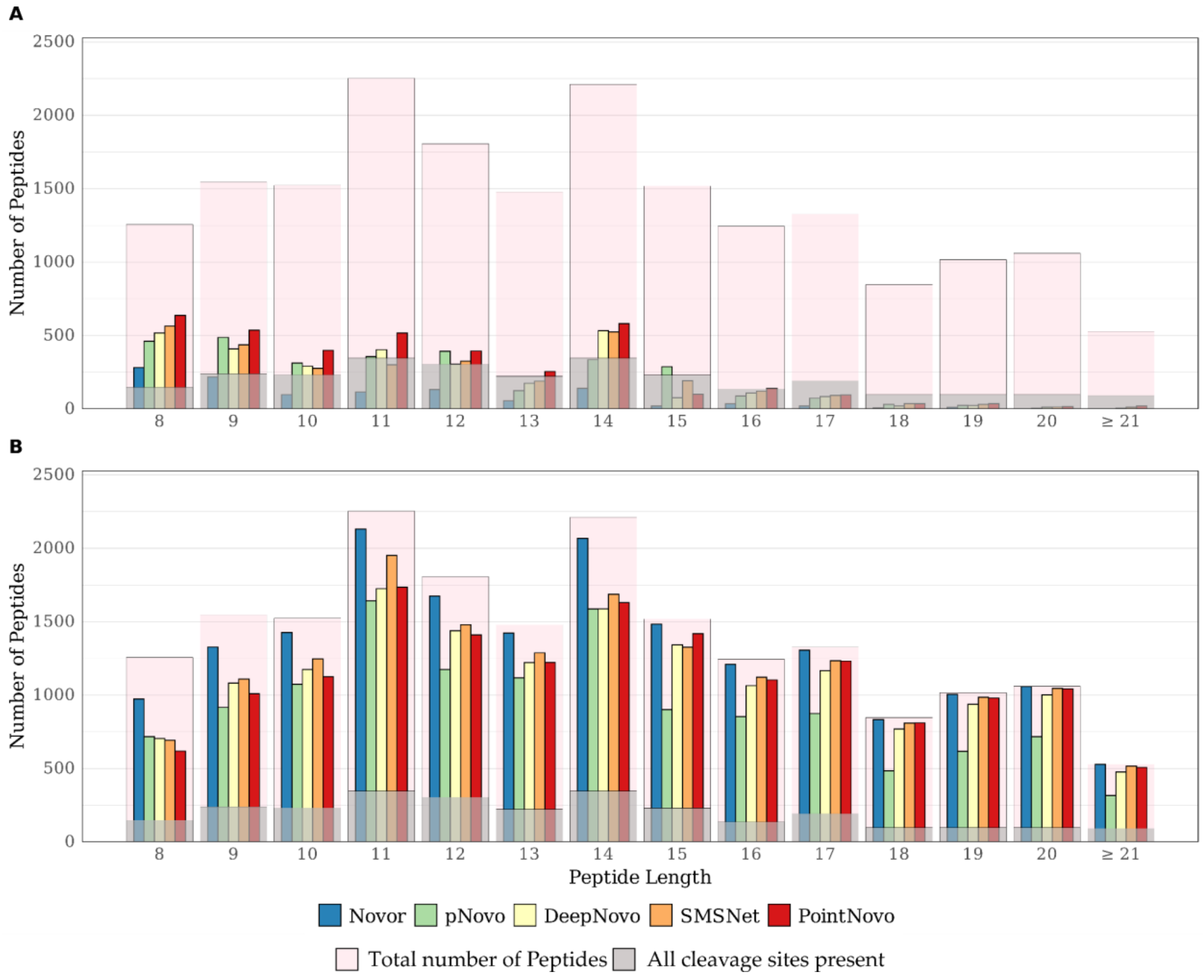


**Supplementary Figure S12** | Peptide length distribution of correct (A) and incorrect (B) predictions of Novor, pNovo 3, DeepNovo, SMSNet and PointNovo on all datasets. The total number of peptides are shown in pink, with the number of peptides without missing cleavage sites shown in gray.

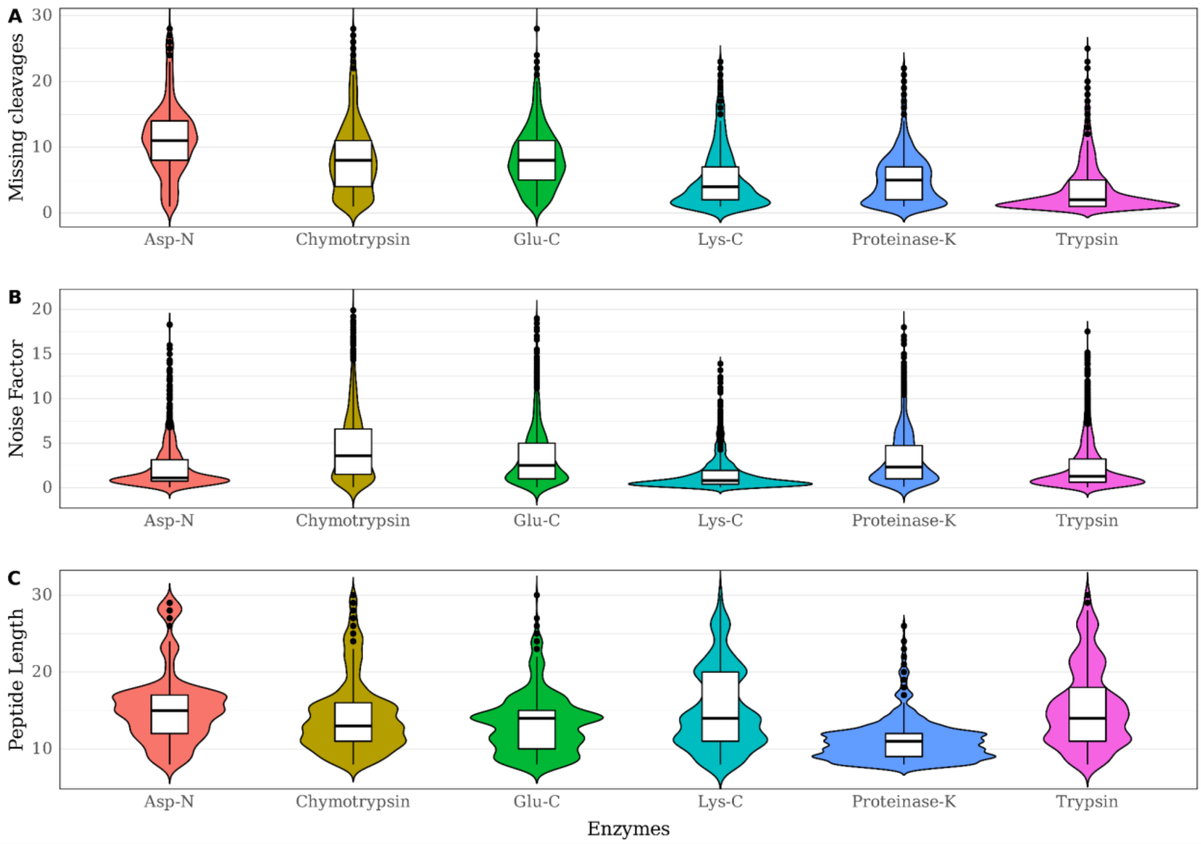


**Supplementary Figure S13** | Violin plots showing the distribution of the number of missing cleavage sites (A), noise factor (B), and length of peptides (C) from different enzymatic datasets of the IgG1-HC. We excluded outliners spectra above a noise of 20 for visualization purposes.

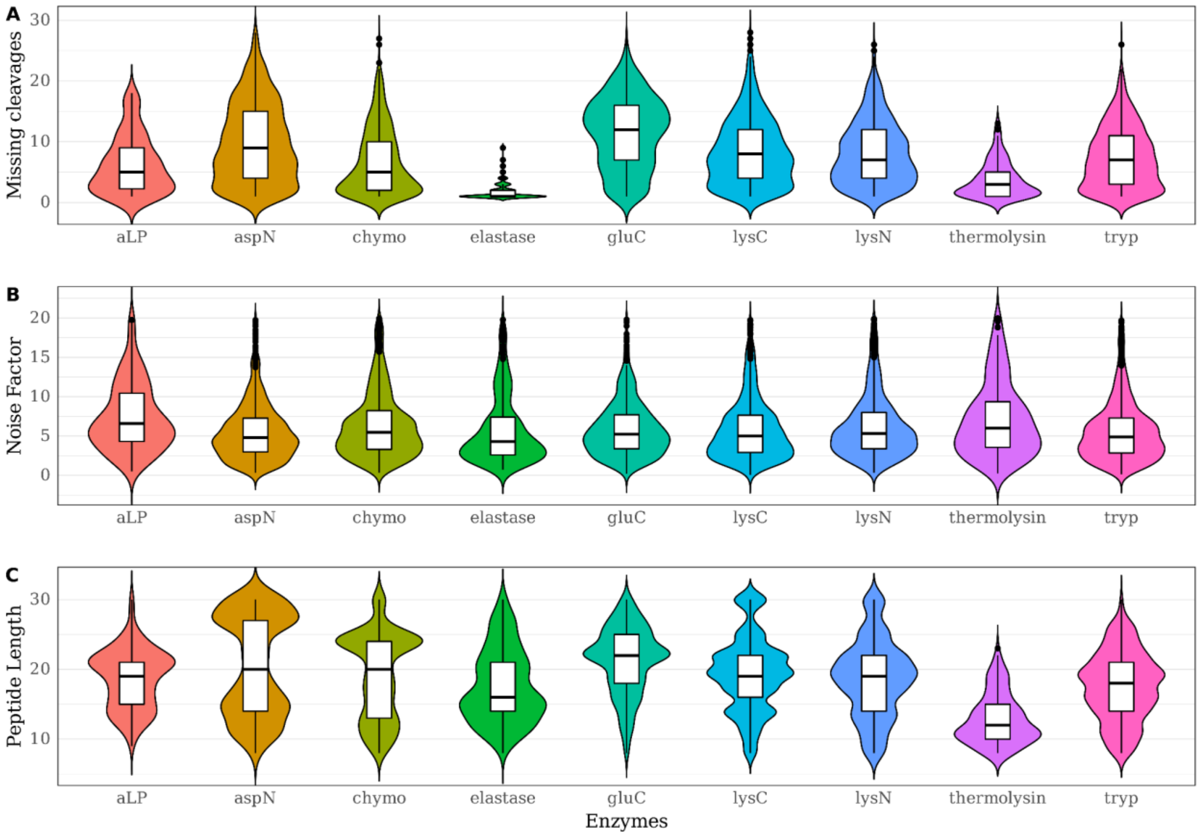


**Supplementary Figure S14** | Violin plots showing the distribution of the number of missing cleavage sites (A), noise factor (B), and length of peptides (C) from different enzymatic datasets of Herceptin. We excluded outliners spectra above a noise of 20 for visualization purposes.

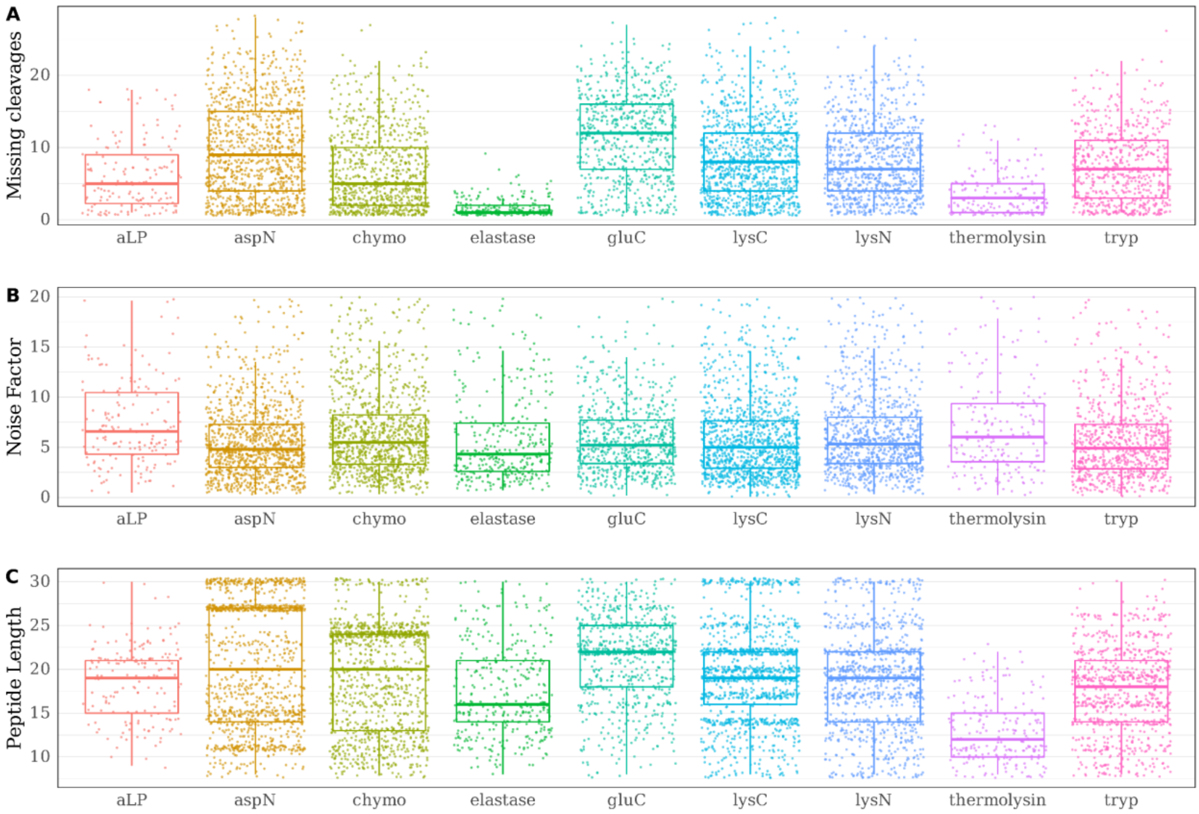


**Supplementary Figure S15** | Scattered boxplots showing the distribution of the number of missing cleavage sites (A), noise factor (B), and length of peptides (C) from different enzymatic datasets of Herceptin. We excluded outliners spectra above a noise of 20 for visualization purposes.

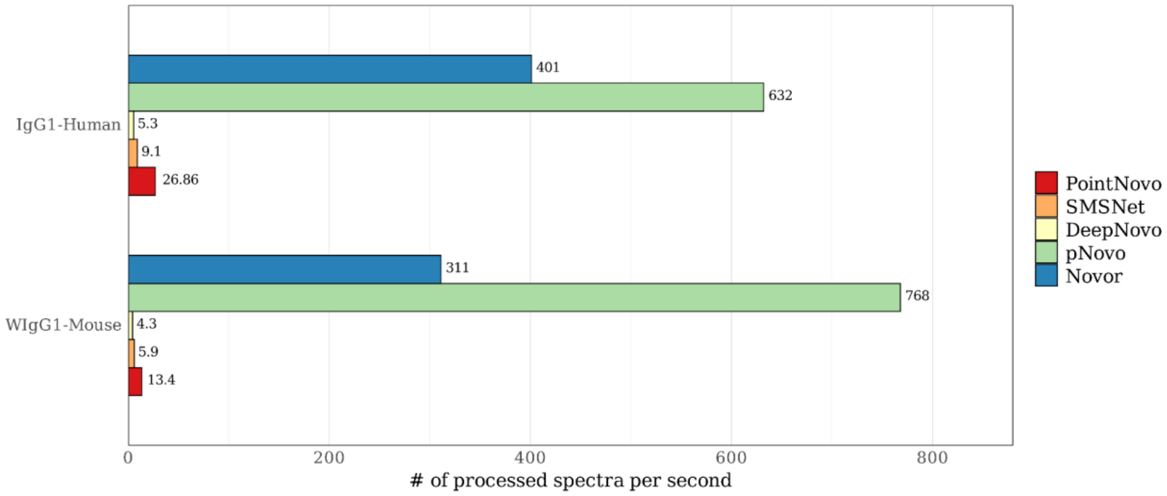


**Supplementary Figure S16** | Average runtime comparison of *de novo* sequencing algorithms. The runtime is shown as the number of spectra processed per second for experimental datasets using Novor, pNovo 3, DeepNovo, SMSNet and PointNovo. For comparison, the average runtime is shown for IgG1-Human (8,735 spectra per file on average) and WIgG1-Mouse (5,858 spectra per file on average).

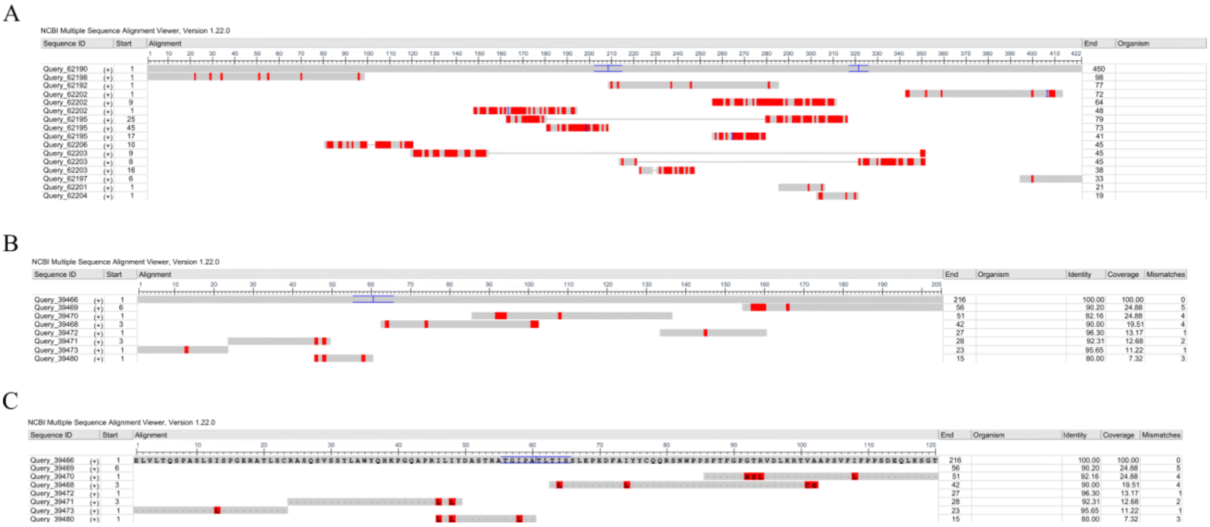


**Supplementary Figure S17** | Assembly results for the IgG1-Human. BLAST alignment of the top assembled contigs from PointNovo against the target heavy chain (A). BLAST alignment of the assembled contigs from PointNovo against the target light chain (B). BLAST alignment of the assembled contigs from PointNovo against the variable region of the target light chain (C).

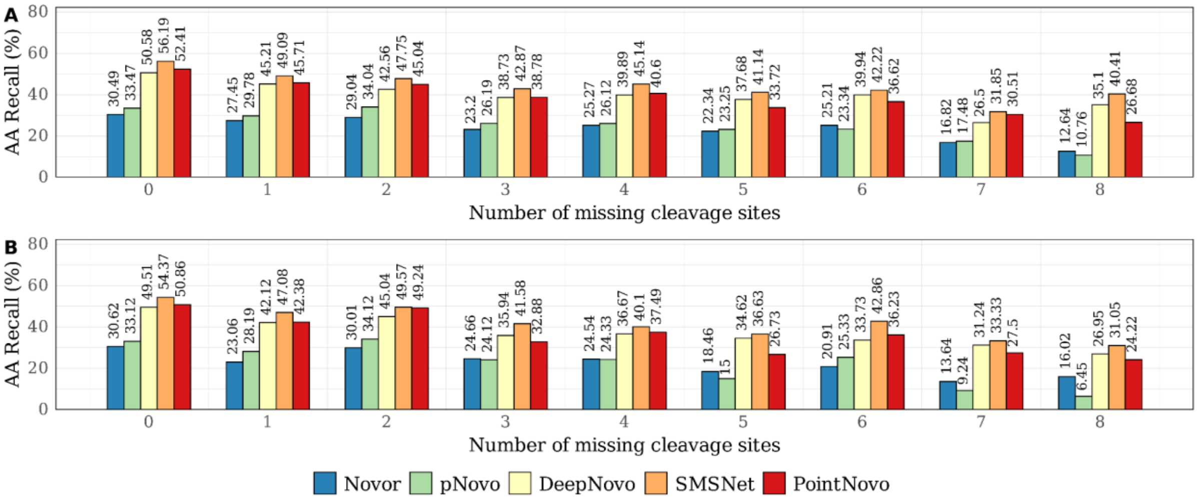


**Supplementary Figure S18** | Total peptide recall of Novor, pNovo 3, DeepNovo, SMSNet and PointNovo across the WIgG1 HC dataset for different number of cleavage sites missing. To evaluate the impact of a-ions, we compared the peptide recall for fragmentation sites with 8 (b and y ions) missing ion types (A) and fragmentation sites with 12 (a, b and y) missing ion types (B).

**Supplementary Table S1** | Incorrect amino acid substitutions made by *de novo* sequencing tools pNovo 3, SMSNet and PointNovo on datasets of IgG1-Human, WIgG1-Mouse, and Herceptin. Shown is the relative amount of specific amino acid substitutions for sequencing errors, where only 1 AA was incorrectly predicted

| Type of Replacement | pNovo 3 | SMSNet | PointNovo |
|---|---|---|---|
| N (deam.) ⇔ D | 59.73 % | 41.64 % | 74.51 % |
| N ⇔ D | 0.84 % | 10.18 % | 7.89 % |
| Q (deam.) ⇔ E | 28.11 % | 7.96 % | 8.86 % |
| K ⇔ Q | 0.00 % | 11.09 % | 0.00 % |
| Other | 11.32 % | 29.13 % | 8.74 % |

**Supplementary Table S2** | Error types made by *de novo* sequencing tools pNovo 3, SMSNet, PointNovo on the datasets of IgG1-Human, WIgG1-Mouse and Herceptin using only peptides without missing cleavages. Shown is the relative amount of 11 different error types for each algorithm.

| Type of Error | pNovo 3 | SMSNet | PointNovo |
|---|---|---|---|
| Number of total predictions | 2203 | 2203 | 2203 |
| Number of total errors | 1589 | 1567 | 1385 |
| Inversion first 3 AAs | 6.30% | 10.12% | 7.53% |
| Inversion last 3 AAs | 2.28% | 4.26% | 4.10% |
| Inversion first & last 3 AAs | 0.00% | 0.09% | 0.11% |
| 1 AA replaced by 1 AA or 2 AAs | 24.51% | 11.26% | 23.74% |
| 2 AAs replaced by 2 AAs | 7.18% | 11.26% | 9.70% |
| 3 AAs replaced by 3 AAs | 5.43% | 5.87% | 7.53% |
| 4 AAs replaced by 4 AAs | 2.01% | 3.97% | 2.96% |
| 5 AAs replaced by 5 AAs | 2.45% | 3.69% | 2.39% |
| 6 AAs replaced by 6 AAs | 4.47% | 1.80% | 4.79% |
| More than 6 AAs wrong | 28.02% | 29.99% | 28.88% |
| Other | 17.33% | 17.69% | 8.21% |

**Supplementary Table S3** | Summary of *de novo* assembly results on heavy chains of three antibody datasets using ALPS (k=7). The top 20 contigs were used to compare the length, coverage, and accuracy of mapped contigs. Mapped contigs must be aligned to the reference protein sequence. The longest contig describes the maximum length of all generated contigs. Sequence coverage was calculated as the percentage of amino acids of the protein sequence that were covered by at least one contig. Accuracy was calculated as the percentage of all protein sequence calls that were correctly labeled.

| | IgG1 HC (446 AA) | WIgG1 HC (441 AA) | Herceptin HC (449 AA) |
|---|---|---|---|
| **pNovo 3** | | | |
| Mapped contigs | 15 | 14 | 14 |
| Longest contig (AA) | 34 (7.62%) | 22 (4.99%) | 42 (9.35%) |
| Sequence coverage (%) | 307 (68.83%) | 211 (47.85%) | 314 (69.93%) |
| Sequence accuracy (%) | 262 (85.34%) | 181 (85.78%) | 242 (77.07%) |
| **SMSNet** | | | |
| Mapped contigs | 13 | 9 | 7 |
| Longest contig (AA) | 65 (14.57%) | 71 (16.10%) | 83 (18.49%) |
| Sequence coverage (%) | 391 (87.67%) | 340 (77.10%) | 314 (69.93%) |
| Sequence accuracy (%) | 348 (89.00%) | 299 (87.94%) | 270 (85.99%) |
| **PointNovo** | | | |
| Mapped contigs | 15 | 10 | 6 |
| Longest contig (AA) | 75 (16.81%) | 66 (14.97%) | 98 (21.82%) |
| Sequence coverage (%) | 437 (97.98%) | 370 (83.90%) | 291 (64.81%) |
| Sequence accuracy (%) | 361 (82.61%) | 347 (93.78%) | 269 (92.44%) |
